# Mechanically Tunable DNA Hydrogel Microparticles for 3D Cellular Systems

**DOI:** 10.1101/2025.07.21.665473

**Authors:** Tobias Walther, Eleni Dalaka, Gotthold Fläschner, Ilia Platzman, Sadaf Pashapour, Michelle Emmert, Pere Roca-Cusachs, Xavier Trepat, Kerstin Göpfrich

## Abstract

Hydrogel microparticles (HMPs) are powerful tools to study and manipulate cellular behavior in 3D cell culture systems and animal models. Here, fully DNA-based HMPs are presented, whose material properties can be precisely tuned by sequence-programmable design of self-assembling DNA nanostructures. These DNA-HMPs offer control over size, stiffness, viscoelasticity and ligand presentation. They are formed by microfluidic encapsulation of two types of orthogonal DNA nanostars and a sequence-complementary DNA linker in water-in-oil droplets. By varying the valency of the DNA nanostar designs, tunable mechanical properties are achieved – spanning three orders of magnitude in Young’s modulus from 30 Pa to 6.5 kPa with distinct viscoelastic behavior. Click-chemistry based functionalization with the small fibronectin-derived peptide cyclic-RGD (c[RGD]) enables integration into fibroblast spheroids. DNA-HMPs are stably retained within the spheroids for several days and undergo design- and stiffness-dependent remodeling, indicating active interactions between the cells and the DNA-HMPs. Combining tunable material properties and inherent biocompatibility of DNA with straightforward functionalization and stimuli-responsiveness, these DNA-HMPs represent a versatile tool to probe and manipulate tissue behaviors in 3D cell cultures and *in vivo* models.

**Table of Contents:** DNA hydrogel microparticles are designed to exhibit controllable viscoelasticity and stiffness across three orders of magnitude from 30 Pa to 6.5 kPa. They are uptaken into fibroblast spheroids where they are actively remodeled by cellular forces depending on their mechanical properties.

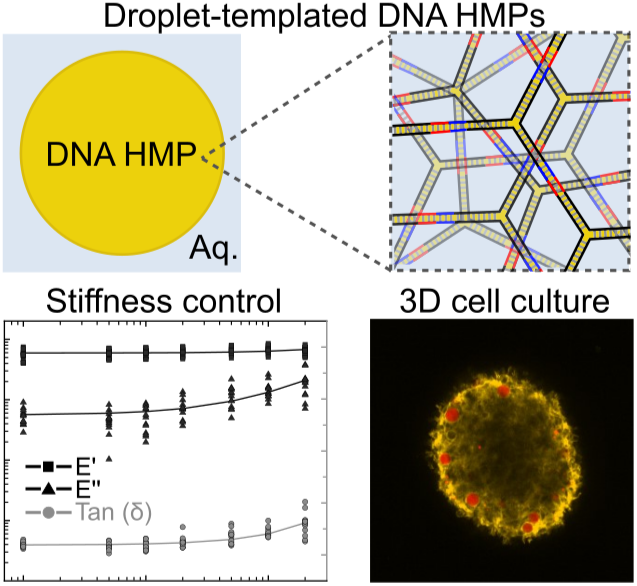

## 1 Introduction

Tools that enable localized perturbation and monitoring of the cell microenvironment are essential to advance 3D cell culture for tissue engineering, disease modeling, and precision medicine [1, 2]. Hydrogel microparticles (HMPs) – micron-sized gel beads – are well-suited for this task. Unlike conventional bulk hydrogels, they are cell-sized objects that allow biochemical and mechanical cues to be applied from within complex multicellular systems, as well as enabling real-time readouts [3, 4, 5, 6]. HMPs have been produced from polymers such as alginate [7, 8, 9], polyacrylamide [10], polyethylene glycol [11], and agarose [12, 13]. They are prominently used as cell culture substrates [14, 15] or as adjuvants that provide physical or chemical signaling [16, 17, 6]. Beyond this, they serve as spatially discrete sensors in tissues owing to their elastic material properties [18, 19, 20, 21]. However, most conventional polymeric HMPs lack versatility in terms of the capacity for targeted chemical functionalization, fine-scale mechanical programmability, and integration of dynamic or stimuli-responsive behavior, such as controlled structural changes or molecular release in response to environmental cues. DNA-based materials offer a promising alternative due to their sequence-defined self-assembly, precise mechanical tuning, and straightforward chemical modification [22, 23, 24]. DNA hydrogels have been applied in various biomedical contexts [25, 26, 27, 28], yet efforts have focused on bulk gels. Despite extensive rheological studies of bulk DNA networks showing storage moduli ranging from 10 Pa to 100 kPa [29, 30, 31, 32, 33, 34], HMPs made from DNA hydrogels with controllable size, mechanical properties, and biofunctionality have not yet been realized. Moreover, such materials have not been explored as deformable force-responsive elements within living tissues. Here, we thus introduce fully DNA-based hydrogel microparticles (DNA-HMPs) with programmable viscoelastic properties and functional biointerfaces. These particles are formed by microfluidic co-encapsulation of self-assembling DNA nanostars and sequence-complementary linkers. They exhibit finely tunable Young’s moduli over three orders of magnitude from 30 Pa to 6.5 kPa that offer exceptional resolution in material design. Beyond microfluidics, DNA-HMPs can also be generated by simple vortexing of the solution making their production broadly accessible without specialized equipment. Once formed, the DNA-HMPs are highly stable and retain their structure for at least 1 year under refrigerated conditions. They behave predominantly elastic under low-frequency indentation, but their viscoelastic response can be modulated through rational DNA nanostar design. The HMPs can be further functionalized via click chemistry with cyclic-RGD (c[RGD])-peptides to enable active cellular interaction. When added to fibroblast spheroids, the DNA-HMPs are readily internalized, stably retained, and undergo design-dependent deformation and remodeling, highlighting their potential as minimally invasive cell-interactive mechanical probes. Combining modular mechanical tuning, biochemical customization, and biocompatibility, DNA-HMPs establish a new class of materials to interact with the 3D cell microenvironment. In the future, DNA-HMPs can therefore serve as a platform for tissue engineering approaches and biophysical analysis in a variety of setups.

## 2 Results and Discussion

### 2.1 Size-controlled formation of DNA-HMPs by microfluidics

Towards applications in 3D cell culture, we first derived a strategy to produce DNA-HMPs in a reproducible and size-controlled manner. We used microfluidics to encapsulate the hydrogel-forming DNA components into water-in-oil droplets. After gel formation, stable DNA-HMPs were released from the droplet shell into an aqueous environment as illustrated in **Figure 1A**. We selected a DNA nanostar design which has been shown to form stable hydrogels in bulk [35, 36]. The design is composed of three single strands of DNA, which form a 3-arm nanostructure upon thermal annealing (for details see Experimental Section). To enable imaging of the resulting DNA-HMPs by fluorescence microscopy, one of the single strands was covalently labeled with a Cyanine-3 (Cy3) dye (for details see Experimental Section). Two sets of such DNA nanostars (A and B), each equipped with orthogonal nine nucleotide-long overhangs on all three arms, form the monomeric units of the hydrogel. As A and B have non-complementary overhangs, cross-linking and hydrogel formation is only initiated upon addition of an 18 nucleotide long single-stranded DNA linker (for DNA sequences see Table S1) [35, 36]. In the following, we will refer to this design as “3-arm short” to distinguish it from other designs that will be introduced later.

**Figure 1:**
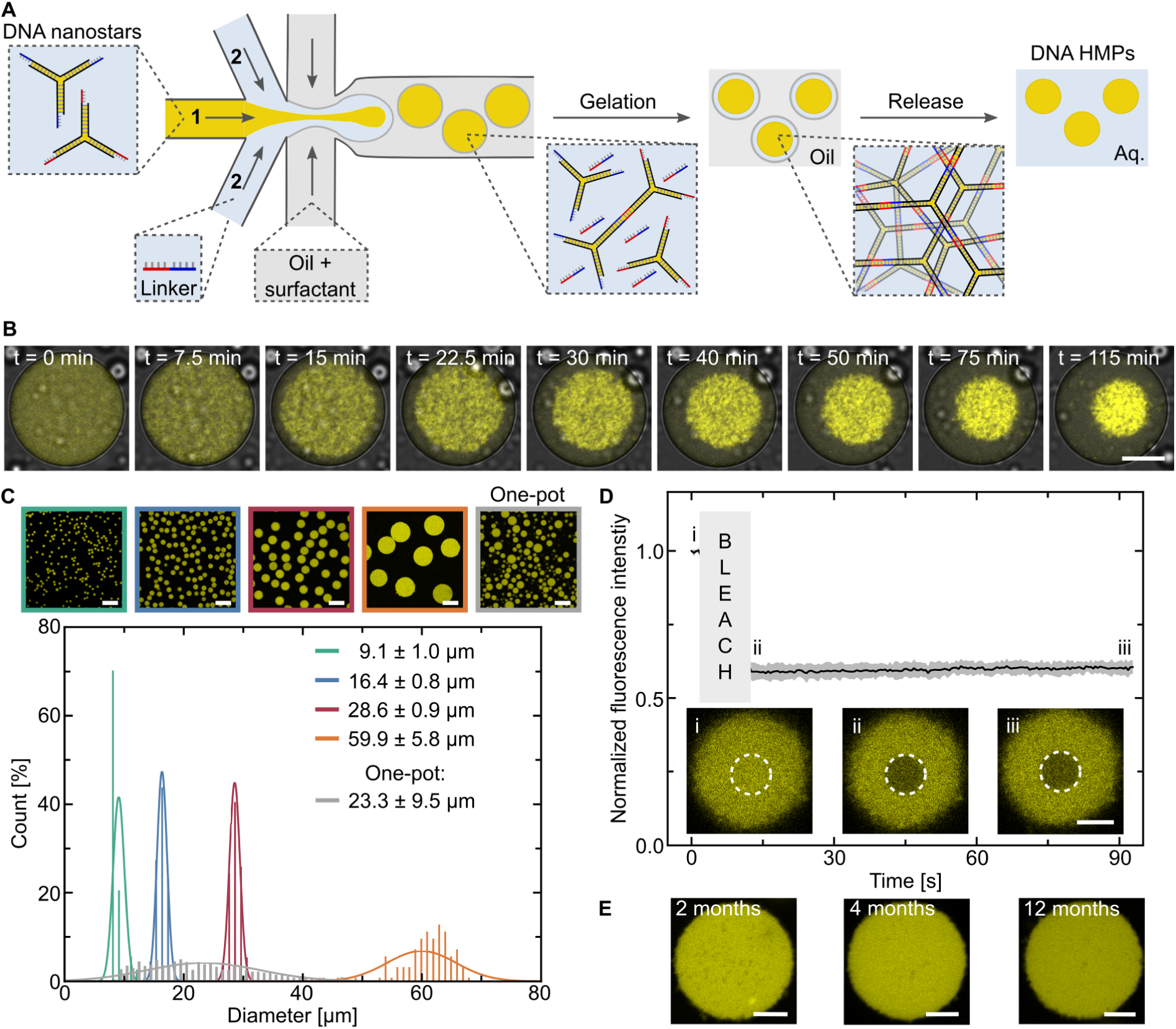
Microfluidic formation of droplet-templated DNA-HMPs. A) Schematic of the procedure for DNA-HMP formation. Two DNA nanostars (A and B) with non-complementary overhangs are encapsulated with a DNA linker using a double-inlet microfluidic device. Upon encapsulation, DNA hydrogel formation is initiated. After full gelation, DNA-HMPs are released from their droplet shell into an aqueous solution. B) Overlay of confocal microscopy (*λ_ex_* = 561 nm, Cy3-labeled DNA, yellow) and brightfield images of a DNA-HMP forming inside a water-in-oil droplet over the course of 115 min, showing how the DNA condenses to form a single DNA-HMP per droplet. Scale bar: 20 µm. C) Confocal microscopy (*λ_ex_* = 561 nm, Cy3-labeled DNA, yellow) images of DNA-HMPs prepared at different monodisperse sizes by controlling the flow rates of the aqueous and oil phases in the microfluidic device. Less uniform DNA-HMPs formed via the one-pot method are shown for comparison (gray distribution). Scale bars: 50 µm. Histograms of the size distributions of the DNA-HMPs with Gaussian fits. n_green_ = 268, n_blue_ = 432, n_red_ = 205, n_orange_ = 125, n_gray_ = 1438. Mean and standard deviation correspond to the mean and standard deviation of the Gaussian fit of each distribution. D) FRAP of DNA-HMPs (mean ± standard deviation, n = 3). Inserted confocal micrographs (*λ_ex_* = 561 nm, Cy3-labeled DNA, yellow) before (*i*) and after bleaching (*ii*, *iii*). Scale bar: 10 µm. E) Stability of DNA-HMPs over time. Confocal microscopy images (*λ_ex_* = 561 nm, Cy3-labeled DNA, yellow) after 2 months, 4 months and 12 months storage in the fridge. Scale bars: 10 µm.

To prevent clogging of the microfluidic device by gel-forming DNA species, we designed a double-inlet microfluidic device where A and B only encounter the DNA linker after the formation of the water-in-oil droplet (for chip layout see Figure S1). To form the DNA-HMPs, A and B were supplied in one inlet, while the second inlet delivered the DNA linker, added at three times molar excess to the 3-arm short DNA nanostars, such that each of the three arms binds one linker. At the intersection of the aqueous phases with a surfactant-oil phase, water-in-oil droplets formed, encapsulating the DNA components.

Sequence-specific binding of the DNA linker to both types of sticky-end overhangs inside the water-in-oil droplets then resulted in the formation of spherical DNA hydrogels which condensed into stable DNA-HMPs over the course of several hours (Figure 1B). Note that for maximum yield, DNA-HMPs were normally matured for 72 h (Figure S2A). The DNA-HMPs were then released from their droplet shells by adding a droplet-destabilizing surfactant, perfluoro-1-octanol, as previously used for the formation and release of droplet-stabilized giant unilamellar vesicles [37]. This allowed us to collect the fully-formed, stable DNA-HMPs in aqueous solution (Figure 1A). Control experiments confirmed that formation of the DNA-HMPs only occurred when both the nanostars and the DNA linker were co-encapsulated (Figure S2B).

Tracking the formation of the DNA-HMPs in water-in-oil droplets over time, we observe the formation of smaller DNA clusters that then condense into a singular spherical DNA-HMP per droplet (Figure S3A, Video S1). During this process, the area fraction of the fluorescent signal inside the droplets and thus the size of the forming DNA-HMP decreases over the course of 2 h before reaching a plateau, indicating maturation of the DNA-HMP (Figure 1B, Figure S3B-D). Cryogenic scanning electron microscopy (cryoSEM) further revealed that the DNA-HMPs consist of interconnected domains of higher DNA density with a diameter on the order of 200 nm (Figure S3E). This architecture indicates that the condensation of the DNA into HMPs is aided by liquid-liquid phase separation on the nanoscale [38].

In order to create uniform DNA-HMPs of varying sizes, we adjusted the flow-rates of the oil- and the aqueous phase, respectively, fine-tuning the diameter of the resulting droplet. We formed DNA-HMPs with narrow size distributions of 9.1 µm ± 1.0 µm, 16.4 µm ± 0.8 µm and 28.6 µm ± 0.9 µm (Figure 1C, for corresponding flow rates see Experimental Section). Utilizing a wider microfluidic channel of 60 µm instead of 30 µm, we were able to produce DNA-HMPs as large as 59.9 µm ± 5.8 µm (Figure 1C), allowing us to mimic a vast range of cell sizes found in mammalian tissues [39]. For the production of large quantities of DNA-HMPs, we additionally developed a scalable one-pot method. It involves layering the DNA-containing solution on top of the oil surfactant mix in a reaction tube or any larger reaction container. Instead of using microfluidics, the formation of water-in-oil droplets is induced by vortexing. While the one-pot method results in a broader size distribution (23.3 µm ± 9.5 µm, Figure 1C), it is fast, scalable and does not require specialized expertise or equipment. It thus facilitates the implementation of DNA-HMPs in laboratories without access to microfluidics when stringent size control is not required. With fluorescence recovery after photobleaching (FRAP) experiments on DNA-HMPs, we confirmed that they consist of a stable gel-phase after release from the droplet shell; the fluorescence of the bleached area did not recover over time (Figure 1D, Video S2). Moreover, the high stability of the DNA-HMPs makes handling straightforward. They can be stored in solution at 4*^◦^*C for at least 1 year without any apparent morphological changes or loss of structural integrity (Figure 1E). Additionally, they settle inside a reaction tube within minutes and can be pelleted by centrifugation using a simple table-top spinner. This allows for quick buffer exchange, washing and functionalization which is key for downstream applications.

### 2.2 Characterization of the material properties of DNA-HMPs tunable by sequence design

Having demonstrated the feasibility of DNA-HMPs, we next explored whether their mechanical properties can be tuned by altering the valency of the DNA nanostars [24, 27, 40, 41] and their arm lengths. Thus, we designed three more DNA nanostars, named 3-arm, 4-arm and 6-arm (Figure 2A). In the updated 3-arm design, the sticky-end overhangs of the 3-arm short DNA nanostar (Figure 2A) were elongated by three additional nucleotides. The DNA linker was extended accordingly, resulting in an increase of the melting temperature from 46*^◦^*C (unbound fraction at 37*^◦^*C = 23%) to 66*^◦^*C (unbound fraction at 37*^◦^*C = 8.8%). We reasoned that this higher thermal stability would improve the suitability of the DNA-HMPs for applications in cell culture and animal models at physiological temperature. Hence, we used the elongated linker also for the higher-valency nanostars with 4 and 6 arms (Figure 2A, for DNA sequences see Table S1).

**Figure 2:**
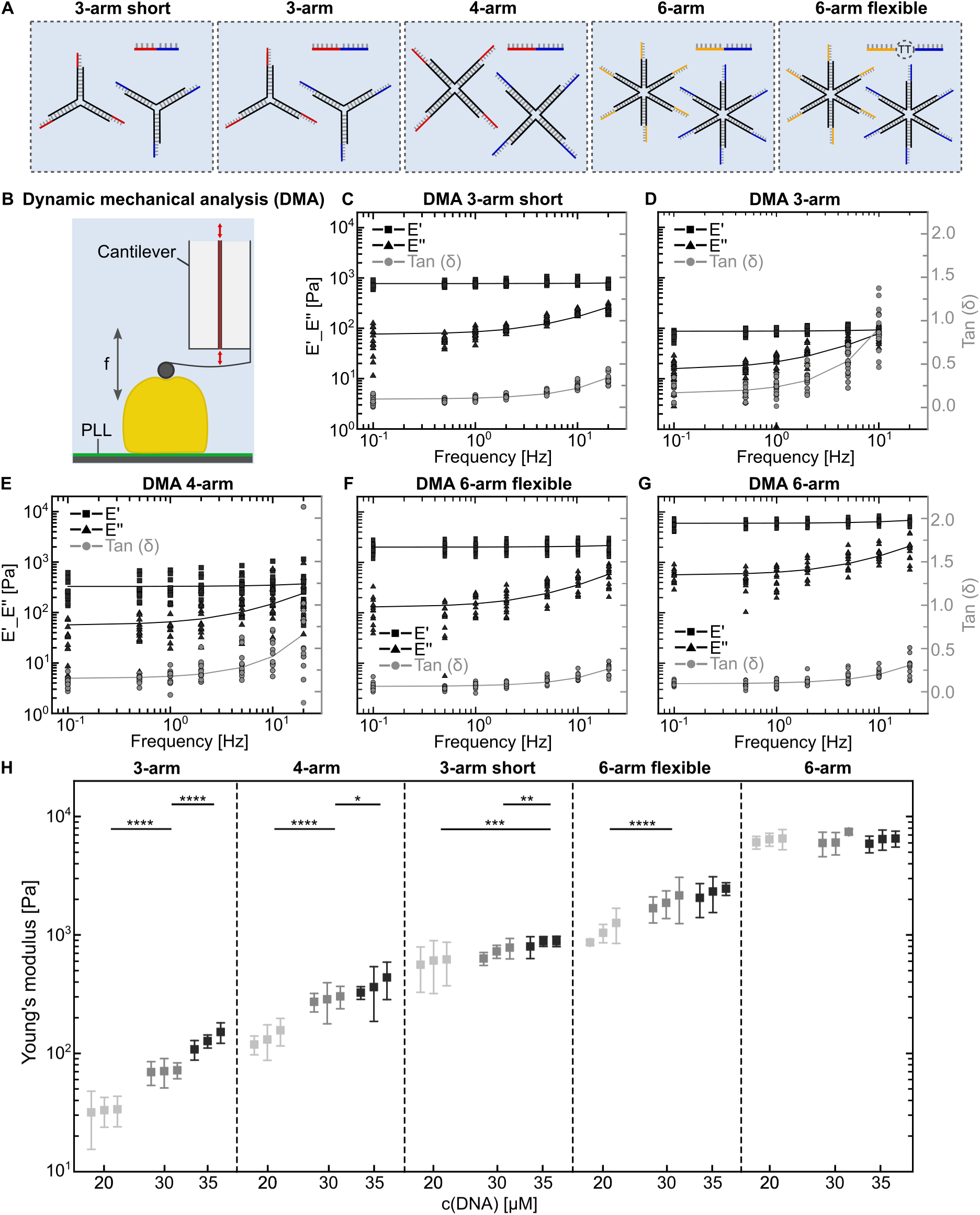
Influence of DNA nanostar sequence design on the material properties of DNA-HMPs. A) Schematic representation of the five different DNA nanostar designs with their respective linkers used to form DNA-HMPs. B) Schematic of HMP indentation using dynamic mechanical analysis (DMA). DNA-HMPs are adhered to a glass substrate by electrostatic interaction with PLL and indented with increaseing frequency *f* using a cantilever. C - G) DMA of 3-arm short (C), 3-arm (D), 4-arm (E), 6-arm flexible (F) and 6-arm (G) DNA-HMPs at 35 µM DNA nanostar concentration. The storage modulus E’ (squares) and the loss modulus E” (triangles) are plotted on a logarithmic scale, tan (*δ*) (E”/E’, filled circles) is plotted on a linear scale. H) Full analysis of Young’s moduli of DNA-HMPs across different designs and DNA concentrations as extracted from Hertz-model fits of the indentation curves during microindentation. The Young’s moduli for 3-arm, 4-arm, 3-arm short, 6-arm flexible and 6-arm DNA-HMPs at DNA monomer concentrations of 20 µM, 30 µM and 35 µM (from left to right) are displayed. The data is presented as mean ± standard deviation of each measurement for n = 3 independent experiments per condition measuring at least 3 particles each. 3-arm ****p-values: 9.9*10^-8^, 3.3*10^-11^. 4-arm *p-value: 0.039, ****p-value: 8*10^-8^. 3-arm short **p-value: 0.003, ***p-value: 0.0008. 6-arm flexible ****p-value: 0.0001. Statistical significance was assessed using unpaired Student’s t-tests.

While DNA-HMPs from the 3-arm and 4-arm DNA nanostars were readily formed at room temperature (Figure S4A/B), 6-arm DNA-HMPs required thermal annealing. This is likely because, partial crosslinking traps the DNA-HMPs in a gel state once three or more arms are bound, while unbound arms act as surfactants, destabilizing the overall structure [42]. Thermal annealing decreases the amount of unbound arms, enabling the formation of stable 6-arm DNA-HMPs (Figure S4C, for details see Experimental Section). To overcome the need for annealing, we reasoned that introducing flexibility in the DNA linkers should also enhance the chances of complete binding. We thus added two flexible thymine-nucleotides to the center of the 6-arm linker (6-arm flexible), which rescued 6-arm DNA-HMP formation at room temperature (Table S1, Figure 2A/S4D). In summary, we developed five distinct DNA nanostar designs that allow for the water-in-oil droplet-templated formation of DNA-HMPs.

Next, we set out to investigate how nanostar valency and design influence the material properties of the DNA-HMPs. Both the stiffness and viscoelasticity of cells and tissues play critical roles in regulating cellular functions [43]. These mechanical properties should therefore be carefully considered when designing materials for tissue engineering applications [44, 45]. Cells exhibit type-specific Young’s moduli ranging from around 0.1 kPa in neurons to 20 kPa and above for osteoblasts [44, 46]. Similarly, the environment and tissues surrounding cells vary greatly from less than 0.1 kPa in mucus up to the GPa range in bone [44, 47]. In addition, cells and their microenvironment also display a broad range of viscoelastic properties [43, 48]. To analyze the different designs and determine their material properties, we turned to microindentation measurements by dynamic mechanical analysis (DMA) as illustrated in Figure 2B. In DMA, a material is exposed to sinusoidal stress as a function of outer parameters such as frequency. Hence, DMA gives insights into the responsiveness of a given material to repeated physical change, allowing for the determination of material properties [49]. For this, the DNA-HMPs were adhered to a glass substrate coated with positively-charged poly-l-lysine (PLL, MW = 150 kDa - 300 kDa) by electrostatic interaction (Figure S5).

3-arm short DNA-HMPs exhibited frequency-independent storage moduli (E’) of 607 Pa ± 287 Pa at 20 µM nanostar concentration to 885 Pa ± 88 Pa at 35 µM nanostar concentration (Figure 2C, Figure S6). In the low frequency range (<1 Hz) the loss moduli (E”) of the DNA-HMPs were substantially lower and likewise frequency-independent. A slight increase of E” was found at higher frequencies (>5 Hz), but never exceeding E’. This is also reflected in the loss tangent *tan*(*δ*), which even at the highest frequency tested (20 Hz) remains <0.5. These data indicate that the 3-arm short HMPs behave largely as elastic materials, displaying viscous dissipation only at high frequencies. These findings are in line with previous work on the rheological behavior of comparable macroscale bulk DNA hydrogels [31, 50, 51, 52].

We next assessed the role of nanostar arm length and valency for the material properties of DNA-HMPs. We hypothesized that elongating DNA nanostar arm lengths would result in a more porous and softer network, owing to the formation of larger nanogel clusters taking up a greater volume fraction of the resulting hydrogel [53, 54]. Conversely, increasing nanostar valency was expected to yield stiffer DNA-HMPs due to higher connectivity within the DNA network [54].

In agreement with these expectations, DMA revealed that the 3-arm DNA-HMPs are softer and more viscous than those formed with the 3-arm short design (Figure 2D, Figure S7). HMPs assembled from the 4-arm design displayed similar viscoelastic behavior at monomer concentrations below 35 µM (Figure S8), while those formed at 35 µM showed increased elasticity (Figure 2E). These results indicate that the viscoelastic properties of our DNA-HMPs can be tuned by nanoscale design. Networks with lower connectivity exhibit more viscous behavior, whereas networks with higher valency and DNA concentration are more elastic. DMA of the 6-arm flexible and 6-arm HMPs further underlined this trend. Across all tested concentrations and frequencies both types of DNA-HMPs showed predominantly elastic behavior with storage moduli reaching up to 2.4 kPa ± 303 Pa for the 6-arm flexible HMPs and 6.5 kPa ± 1 kPa for the 6-arm HMPs (Figure 2F/G, Figure S9/S10). Comparing these findings with literature, we find that they are well in line with reports showing lower elastic moduli in macroscale bulk DNA hydrogels with increased flexibility [31, 51, 55]. Consistent with DMA analysis, the relaxation behavior of the DNA-HMPs following indentation showed a more viscous response for the 3-arm and 4-arm designs compared to the other HMPs (Figure S11). Indeed, 3-arm and 4-arm HMPs showed stronger decrease in initial load prior to relaxation, indicating higher viscosity, while both types of 6-arm HMPs exhibited less load decrease and thus a more elastic response (Figure S11). Finally, increasing the DNA monomer concentration also increased DNA-HMP stiffness, albeit to a lesser extent than sequence design (Figure 2H). Taken together, our findings indicate that DNA-HMPs function as microscale DNA hydrogels whose mechanical properties can be precisely modulated within the cell stiffness and viscoelasticity range, exhibiting cell-like loss tangent behavior [56], by adjusting nanostructural design.

### 2.3 Real-time deformability cytometry of DNA-HMPs

To examine the material properties of DNA-HMPs at high-frequency manipulation and to verify their Young’s moduli with an independent measurement, we subjected our DNA-HMPs to real-time deformability cytometry (RT-DC); a microfluidic method used to measure cell stiffness in a contact-free and high- throughput manner [57]. Cells are flushed through a narrow channel such that they deform under hydrodynamic shear force. From the deformation and flow speed, it is possible to infer the Young’s modulus of thousands of cells within minutes [57, 58, 59].

The DNA-HMPs deformed upon entering the RT-DC channel, relaxing into a steady-state deformation once fully inside the channel. After leaving the channel, they returned to their initial spherical shape, indicating the absence of plastic deformation under the applied shear force (Figure 3A). The images presented in Figure 3A were acquired by increasing the size of the imaged channel region during RT-DC to span the entire length of the channel and beyond, while increasing the imaging speed to 7000 f/s, tracking HMP deformation in a quasi-dynamic RT-DC (dRT-DC) experiment [59]. Given the low optical contrast of our HMPs, we adjusted the RT-DC workflow to include a 20x phase-contrast objective (see Experimental Section). Comparison of the deformation during RT-DC for the 3-arm short, 6-arm flexible, and 6-arm DNA-HMPs reveals clear design-dependent differences in deformability. The 3-arm short HMPs show the highest deformation adopting a bullet shape as they transverse the channel, while deformation is less pronounced for the 6-arm flexible design and nearly absent for 6-arm HMPs (deformation ≤ 0.03, Figure 3A/B).

**Figure 3:**
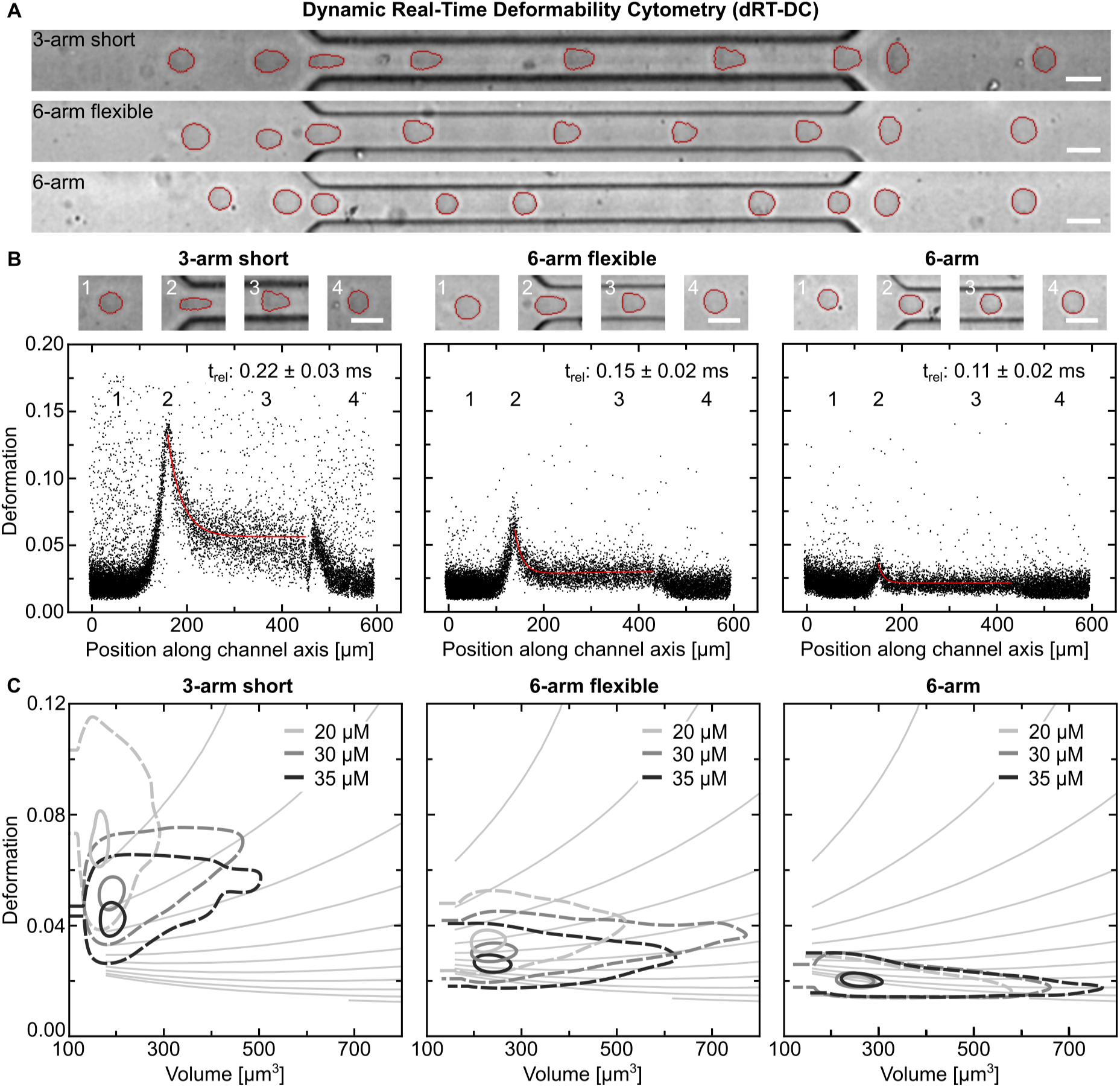
Analysis of the material properties of DNA-HMPs using real-time deformability cytometry (RT-DC). A) Composite images of DNA-HMPs (3-arm short, 6-arm flexible and 6-arm) inside the flow channel during dynamic RT-DC (dRT-DC). DNA-HMPs are initially spherical and deform upon entering the channel. Within milliseconds, they reach a steady-state deformation. After leaving the channel, they return to their spherical shape. Scale bar: 20 µm. B) Deformation scatter plots of 35 µM 3-arm short, 6-arm flexible and 6-arm HMPs plotted over the channel length during dRT-DC. Each point corresponds to an individual DNA-HMP (n = 13000). Insets 1 - 4 show DNA-HMPs at the corresponding positions during dRT-DC depicted in the scatter plots. The relaxation time t_rel_ of the DNA-HMPs was extracted from the exponential fit of the relaxation scatter plot (red curve). For calculation of the flow time see Experimental Section. Scale bars: 20 µm. C) Exemplary contour plots showing DNA-HMP deformation over the particle volume for different DNA nanostar concentrations for the 3-arm short, 6-arm flexible and 6-arm designs. The data is presented using contour plots showing the 50^th^ percentile (dashed line) and 95^th^ percentile (solid line). Isoelasticity lines derived from numerical simulations are shown additionally, indicating stiffness changes where a steeper slope corresponds to softer particles. Only one repeat per condition is depicted to improve readability. For the full set of replicates see Figure S12.

As shown in Figure 3B, DNA-HMP deformation peaked for all three designs just after entering the channel and rapidly decreased during relaxation into a steady-state followed by a return to a spherical shape upon leaving the channel. The differences between the three designs can be seen from the marked decrease in initial deformation from 0.15 for the 3-arm short HMPs to 0.075 for the 6-arm flexible design and 0.03 for the 6-arm HMPs, which thus remained almost spherical inside the channel, exhibiting a weak bullet shape. Exponential fits to the slope of DNA-HMP relaxation following initial deformation revealed characteristic response times *τ* of 0.22 ± 0.03 ms (3-arm short), 0.15 ± 0.02 ms (6-arm flexible) and 0.11 ± 0.02 ms (6-arm) respectively (Figure 3B), showing faster relaxation responses for higher valency, stiffer DNA-HMPs.

Plotting DNA-HMP deformation against their volume, we show that an increase in DNA nanostar concentration also resulted in a decrease in deformability, in line with our results obtained by DMA (Figure 3C). As already seen during dRT-DC measurements, the 6-arm HMPs did not exhibit strong deformation and in fact remained almost spherical with all conditions showing a mean deformation of around 0.02 (full data presented in Figure S12). Based on numerical simulations for fully elastic spheres, RT-DC is normally used to determine apparent Young’s moduli from such deformation data [57, 58]. As the DMA data suggests that our DNA-HMPs behave as viscoelastic solids, we can also extracted their Young’s moduli from the RT-DC. We find that Young’s moduli determined with RT-DC are in good agreement with the microindentation data for the 3-arm short and 6-arm flexible HMPs (Figure S13/S14). Given the low deformation of the 6-arm HMPs during RT-DC at the tested flow rate of 0.04 µL*/*s, accurate stiffness determination is not possible for this design. Nonetheless, the data gathered via RT-DC shows the same trend as the microindentation (Figure S15).

While DMA by microindentation allowed us to investigate the rheological properties of the DNA-HMPs on the second time scale, RT-DC allows us to study relaxation on the millisecond time scale. Based on both the response time *τ*, extracted by fitting the dynamic RT-DC data, and Young’s modulus *E* we were able to calculate apparent viscosities *η* of the DNA-HMPs following *τ* = *η/E* as 0.167 Pa s (3-arm short), 0.211 Pa s (6-arm flexible) and 0.311 Pa s (6-arm) [59]. DNA-HMPs thus show a mostly elastic response to the applied forces exhibiting low, but measurable, viscosity.

Similar analysis of the DNA-HMPs at 20 µM DNA nanostar concentration revealed slower relaxation times and thus a decreased elasticity, well in line with our data gathered by DMA (Figures S16-S18).

We were not able to fit the shape of the soft DNA-HMPs of the 3-arm and 4-arm designs accurately owing to their low optical contrast. Similar findings have been reported for polyacrylamide hydrogel microparticles below a stiffness of 700 Pa [19].

### 2.4 Biofunctionalization of DNA-HMPs and design-specific mechanical sensing in a 3D cellular system

Finally, we examined whether DNA-HMPs can serve as mechanical sensors in 3D cellular systems. We integrated DNA-HMPs of the 3-arm, 4-arm, 6-arm flexible and 6-arm designs into 3D fibroblast spheroids. For this, mouse liver fibroblasts expressing td-Tomato were co-cultured for 48 h with DNA-HMPs in hanging drops [60] (Figure 4A, Videos S3-S6). DNA-HMP uptake by the spheroids was facilitated by addition of the short fibronectin-derived peptide sequence c[RGD] to the DNA linkers. c[RGD]-tagged DNA linkers were created using Dibenzocyclooctin (DBCO)-azide click chemistry of DBCO-tagged DNA single strands (see Table S1) and an azide-modified c[RGD] moiety. Formation of the c[RGD]-tagged DNA linkers was confirmed using polayacrylamide gel electrophoresis (PAGE, Figure S19). Spheroids developed normally in the presence of DNA-HMPs, which were stable inside the spheroids during the 48 h growth period. The DNA-HMPs were not expelled from the spheroids during the subsequent observation time, showing their stability in 3D cell culture.

**Figure 4:**
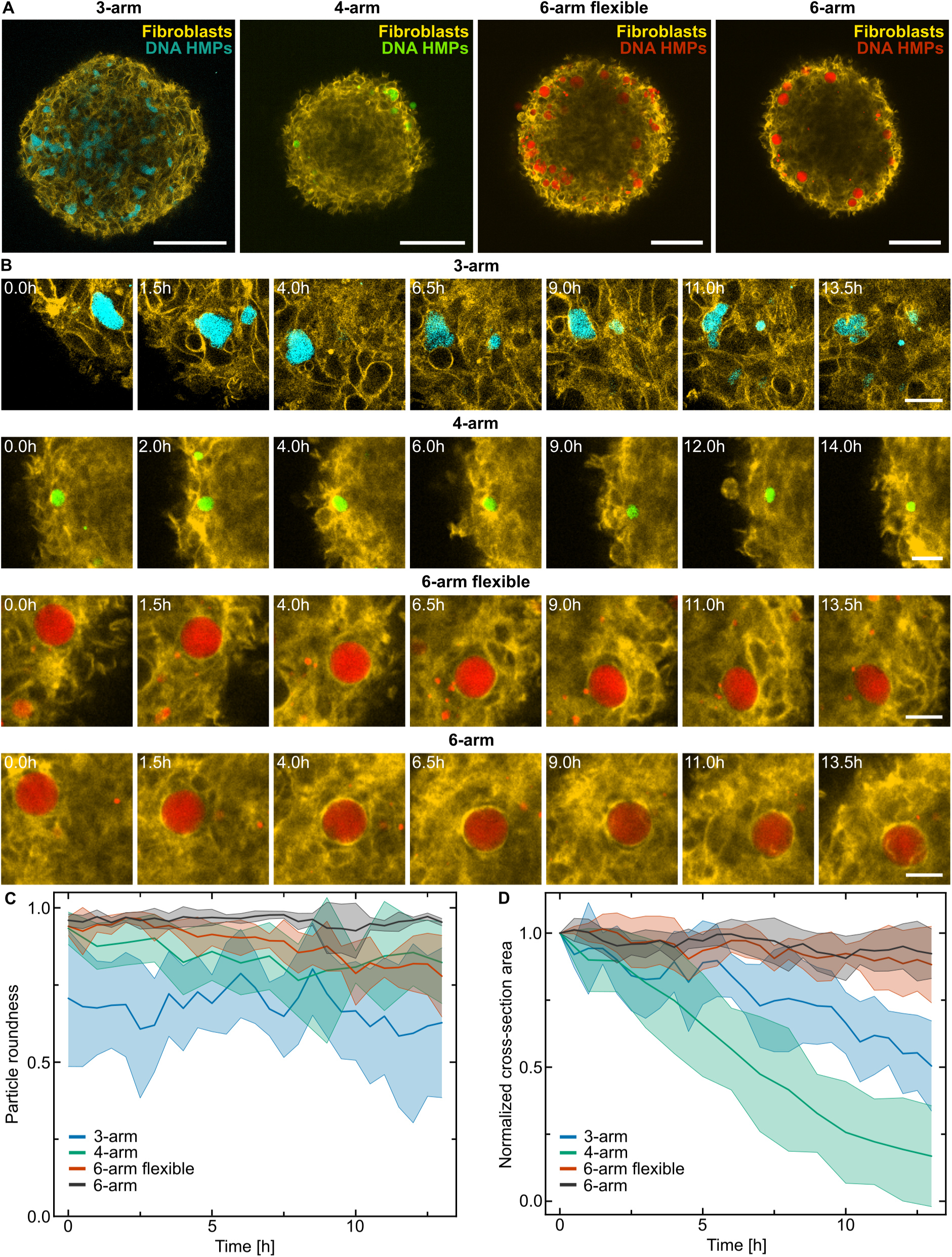
Integration of DNA-HMPs into 3D fibroblast spheroids. A) Confocal microscopy of fibroblast spheroids (*λ_ex_* = 561 nm, td-Tomato, yellow) with embedded 3-arm (*λ_ex_* = 405 nm, ATTO-390-labeled DNA, cyan), 4-arm (*λ_ex_* = 488 nm, ATTO-488-labeled DNA, green), 6-arm flexible (*λ_ex_* = 640 nm, ATTO-647N-labeled DNA, red) and 6-arm (*λ_ex_* = 640 nm, ATTO-647N-labeled DNA, red) DNA-HMPs after 48 h of hanging drop co-culture. Scale bars: 100 µm. B) Equatorial z-slices of 3-arm, 4-arm, 6-arm flexible and 6-arm DNA-HMPs embedded in 3D fibroblast spheroids showing particle deformation observed over time. Scale bars: 20 µm. C) Particle roundness of DNA-HMPs inside 3D fibroblast spheroids extracted from 2D z-slices plotted over time. D) Cross-sectional area of DNA-HMP equatorial planes extracted from 2D z-slices over time, normalized to the first time-point. The data in C and D is presented as mean ± standard deviation per DNA nanostar design measuring five DNA-HMPs each.

Anaylsis of 2D z-slices of the DNA-HMPs during continuous observation for several hours revealed that DNA-HMPs are deformed by the surrounding mouse fibroblasts in a design-dependent manner. While 3-arm HMPs are strongly deformed, 4-arm HMPs remain more spherical, but are still compressed over time. The 6-arm flexible and 6-arm HMPs retained a more spherical shape, with the stiffest design showing minimal deformation (Figure 4B, Videos S7-S10). Analysis of the roundness (Figure 4C) and the compression (Figure 4D) of individual DNA-HMPs inside the spheroids is consistent with their respective rheological properties. The softer DNA-HMPs decrease their cross-sectional area with time, indicating that an increasing compressive stress generated by fibroblasts is sufficient to compact the HMPs. By contrast, the stiffer 6-arm DNA-HMPs maintain the same cross-sectional area over the 14 h observation period (Figure 4D). The strong decrease in size of the 4-arm HMPs compared to the 3-arm design may be explained by the difference in viscosity between the two designs. As the 3-arm HMPs are more viscous in nature, they seem to show rather complex remodeling instead of the plastic compaction observed for the 4-arm HMPs. Due to their soft nature, the 4-arm HMPs get compacted rather than sheared like the 3-arm design, which can also explain their relative roundness over time.

## 3 Conclusion

In this work, we show the formation and characterization of DNA-HMPs for 3D cell culture applications. We demonstrate that their size, stiffness, viscoelasticity and functionalization are controllable by sequence design alone; a property which is difficult to achieve to the same extent with conventional hydrogel systems. In particular, the Young’s modulus of our DNA-HMPs is finely tunable across three orders of magnitude from 30 Pa to 6.5 kPa. While low-valency 3-and 4-arm DNA nanostar designs result in very soft and more viscous HMPs, higher valency 6-arm HMPs exhibit stiffer and more elastic behavior. Intriguingly, this switch is tunable by changing the DNA overhang design, as shorter overhang 3-arm structures are more elastic and stiffer than those with longer overhangs. This suggests that with changes as small as 3 nucleotides, DNA hydrogel network properties can be fundamentally altered. This property warrants further exploration and could lead to evolvable “Darwinian” materials. Finally, we show that c[RGD]-functionalized DNA-HMPs are uptaken into 3D spheroid structures. Further, depending on the material properties of the spheroid-embedded HMPs, the surrounding fibroblasts are able to interact with and deform them.

The DNA-HMPs are delivered into 3D cellular systems in a completely non-invasive manner thanks to the functionalization with cell-binding peptides, opening up the possibility for their further usage in other 3D or 2D multicellular systems. Combined with the already established stimuli-response system for the release of cargo [6], such DNA-HMPs could offer a unique multi-functionality whereby the particles allow for locally-confined chemical perturbations in combination with a mechanical readout. Given the ease of design, the self-assembly of DNA hydrogel structures and the myriad of possible stimuli-responses such as temperature, light, pH or protein interaction inherent to DNA, DNA-HMPs could become powerful tools for the further study and manipulation of engineered cellular systems.

## 4 Experimental Section

### Design and handling of DNA sequences

The design and sequence of the DNA strands used for 3-arm short nanostars A and B as well as the linker strand were adapted from previous publications [35, 36]. The sequences for the 3-arm nanostars C and D were designed based on these by elongating the linker and the respective sticky-end overhangs by three additional nucleotides each.

The 4-arm nanostars were based on the nanostar design from a previous publication [40] and adapted to contain the sticky-end overhangs for the elongated linker. Likewise, the 6-arm nanostars were based on another structure from a previous publication [41], adapted to contain sticky-end overhangs and a corresponding linker of the same length as the elongated linker. Melting curves and binding behavior of all structures were analyzed and verified using NuPack [61]. All DNA was purchased either from Integrated DNA Technologies (unmodified DNA, purification: standard desalting) or Biomers (modified DNA, purification: HPLC). All unmodified DNA was diluted in 10 mM Tris (pH = 8) and 1 mM EDTA (Sigma Life Science) to yield 800 µM stock solutions. Modified strands were diluted in MilliQ water to yield 800 µM stock solutions. DNA concentrations were measured using a nanodrop device (Implen). All DNA sequences are listed in Table S1, Supporting Information. The DNA stock solutions were stored at −20*^◦^*C, if not in use.

### Preparation of 3-arm short and 3-arm DNA nanostars

Annealing of the DNA nanostars (A/B and C/D) was achieved by mixing the three respective DNA single-strands A-1, A-2, A-3 or B-1, B-2, B-3; C-1, C-2, C-3 or D-1, D-2, D-3 at equimolar ratios resulting in 150 µM solutions of the DNA nanostars A and B or C and D. For confocal fluorescence microscopy, 4 mol% of cyanine-3 (Cy3)-labeled strands (B-1-Cy3 and D-1-Cy3) or Atto-390-labeled strands (D-1-390) were added. Phosphate-buffered saline (PBS, no CaCl_2_, no MgCl_2_, pH = 7.4; Gibco) solution at a final concentration of 1x, as well as a final concentration of 10 mM of MgCl_2_ were added for annealing. The nanostars were annealed in a thermal cycler (BioRad) by heating the samples to 85*^◦^*C for 3 min and subsequently cooling to 20*^◦^*C at an increment rate of 0.1*^◦^*C/s. If not stated otherwise, 1x PBS and 10 mM MgCl_2_ were used as buffer conditions for all experiments.

### Preparation of 4-arm DNA nanostars

Annealing of the 4-arm DNA nanostars (E/F) was achieved by mixing the four respective DNA single-strands E-1, E-2, E-3 and E-4 or F-1, F-2, F-3 and F-4 at equimolar ratios resulting in 150 µM solutions of the 4-arm design. For confocal fluorescence microscopy, 4 mol% of an Atto-488-labeled DNA strand (E-1-488) were added to the E monomer solution. Phosphate-buffered saline (PBS, no CaCl_2_, no MgCl_2_, pH = 7.4; Gibco) solution at a final concentration of 1x, as well as a final concentration of 10 mM of MgCl_2_ were added for annealing. The 4-arm design was annealed in a thermal cycler (BioRad) by heating the samples to 95*^◦^*C for 10 min and subsequently cooling to 20*^◦^*C at an increment rate of 0.6*^◦^*C/min.

### Preparation of 6-arm DNA nanostars

Annealing of the 6-arm DNA nanostars (G/H) was achieved by mixing the six respective DNA single-strands G-1, G-2, G-3, G-4, G-5, and G-6 or H-1, H-2, H-3, H-4, H-5 and H-6 at equimolar ratios resulting in 117 µM solutions of the 6-arm design. For confocal fluorescence microscopy, 4 mol% of an Atto-647N-labeled DNA strand (G-1-647) were added to the G monomer solution. Phosphate-buffered saline (PBS, no CaCl_2_, no MgCl_2_, pH = 7.4; Gibco) solution at a final concentration of 1x, as well as a final concentration of 10 mM of MgCl_2_ were added for annealing. The 6-arm design was annealed in a thermal cycler (BioRad) by heating the samples to 95*^◦^*C for 10 min and subsequently cooling to 20*^◦^*C at an increment rate of 0.6*^◦^*C/min.

### Preparation of microfluidic chips

All microfluidic chips were designed using the CAD software QCAD-pro (Ribbonsoft, Switzerland). The channels are 30 µm (standard) or 60 µm (only for the largest beads in Fig. 1) in height and width respectively. SU8-3025 negative photoresist (MicroChem, USA) was spin-coated (Laurell Technologies Corp., USA) at 2600 rpm for 30 s to achieve a 30 µm uniform layer in height on 2-inch silicon wafers (MicroChemicals, Germany), twice that for the 60 µm channel. The designed structures were then exposed to the photoresist-coated wafer by using the Tabletop Micro Pattern Generator µMLA (Heidelberg Instruments, Germany) with 375 mJ*/*cm^2^. The wafer was baked at 65*^◦^*C for 1 min and for another 5 min at 95*^◦^*C to harden the exposed regions and developed afterwards to remove non-exposed photoresist with a developer (mr-DEV 600, MicroChemicals, Germany). A hard bake was performed at 150*^◦^*C for 30 min. Microfluidic chips were produced using soft lithography. Briefly, Polydimethylsiloxane (PDMS, Sylgard 184, Dow Corning, USA) was prepared in a 9:1 (w/w) ratio (oligomer:polymerization catalyst). The solution was used to cover the wafer and hardened at 65*^◦^*C for at least 2 h following degassing of the PDMS-solution under vacuum. The PDMS block was then cut out, and connection holes for the inlets and outlets of 0.5 mm size (Biopsy Punch, World Precision Instruments, USA) punched. As a final step the PDMS block was cleaned with 70% EtOH and activated in an oxygen plasma (300 Semi-auto plasma processor (PVA TePla AG 0.5 mbar, 200 W, 30 s)) together with a coverslip (Carl Roth, Germany, 24 x 60 mm) and bonded. The bonded chips were incubated at 65*^◦^*C overnight and stored at room temperature until further usage. Prior to DNA-HMP production, the microfluidic channels were coated using a perfluorinated oil (Aquapel).

### Preparation of droplet-templated DNA-HMPs using the 3-arm short design

DNA-HMPs were produced in a water-in-oil droplet-templated manner following encapsulation of the DNA-containing solution into water-in-oil droplets. Water-in-oil droplets were either produced by microfluidics (for the data shown in Figure 1D) to ensure uniform droplet sizes, or by manual shaking (for all other presented data).

For water-in-oil droplet production in a microfluidic setup, channels containing a flow-focusing T-junction were used. For DNA-HMPs of up to 30 µm in size, a double-inlet microfluidic chip was used (channel design enclosed as Figure S1A, Supporting Information). Here, the nanostars A and B were supplied at equimolar concentrations (20 µM each) in one channel, while the linker strand was supplied through the second inlet at a 3x higher concentration to ensure gel-formation (60 µM linker strand for 20 µM DNA nanostars). The DNA solutions were supplied using a syringe pump system (NemeSys, Cetoni). Mixing of the two phases and thus gel-formation was facilitated upon encapsulation into water-in-oil droplets at the water-oil junction. To ensure equal mixing of both nanostars and linker strands, both aqueous streams were injected using the same flow rates. As the oil-phase, a solution of 2 wt% perfluoropolyether–polyethylene glycol (PFPE–PEG, RAN Biotechnologies) dissolved in HFE-7500 (Iolitex Ionic Liquids Technologies) was supplied using a separate syringe pump (World Precision Instruments) at a ratio of 4:1 oil to aqueous flow rates (e.g. 0.4 µL*/*s oil:0.1 µL*/*s aqueous phase). For DNA-HMPs smaller than 30 µm, up to three times higher oil flow rates were applied. DNA-HMPs larger than 30 µm were produced in a single-inlet microfluidics device (channel design enclosed as Figure S1B) using an air-pressure-based elveflow control microfluidics system (Elveflow). Here, the linker strands were added to the nanostar-containing solution directly before applying it to the device to keep the gelation time outside of the droplets minimal. The aqueous and oil solutions were supplied in a ratio of 300 mbar to 200 mbar of the PFPE-PEG containing oil phase and the sample-containing aqueous phase, respectively.

Alternatively, water-in-oil-droplets containing the same oil and aqueous phase were created by adding the aqueous solution on top of the oil phase in a volumetric ratio of 1:3 in a microtube (Eppendorf, typically 50 µL aqueous to 150 µL oil phase) and flicking the tube with a finger 8 times [37].

The droplets were then stored at 22*^◦^*C room temperature for 72 hours to allow the DNA-HMPs inside the droplets to fully assemble. After that, DNA-HMPs were released from the water-in-oil emulsion by first adding the buffer in which the DNA-HMPs were formed (always: 1x PBS and 10 mM MgCl_2_) at 1.5x the initial volume on top of the droplet emulsion and subsequently breaking the emulsion. To break the droplet emulsion, 1H,1H,2H,2H-Perfluoro-1-octanol (Merck) was added on top of the buffer and droplet emulsion, and incubated for 30 min. Released DNA-HMPs in solution were then transferred to a new microtube for use.

### Preparation of 3-arm, 4-arm and 6-arm flexible DNA-HMPs

3-arm, 4-arm and 6-arm flexible DNA-HMPs were created similarly to the 3-arm short DNA-HMPs following the same process of templated formation of DNA-HMPs in water-in-oil-droplets. Here, only manual shaking was used to form the DNA-HMPs presented in this study.

For the formation of 3-arm DNA-HMPs, the 3-arm nanostars C and D were mixed at equimolar ratios (e.g. 20 µM each) and the addition of 3x this concentration of the elongated linker (e.g. 60 µM for 20 µM DNA-HMPs).

For the formation of 4-arm DNA-HMPs, the 4-arm nanostars E and F were mixed at equimolar ratios (e.g. 20 µM each) and the addition of 4x this concentration of the elongated linker (e.g. 80 µM for 20 µM DNA-HMPs).

Likewise, 6-arm flexible DNA-HMPs were prepared following the same experimental setup, but using equimolar concentrations of the nanostars G and H and 6x the corresponding concentration of the flexible 6-arm-linker (e.g. 120 µM for 20 µM DNA-HMPs). To achieve even gelation, monomer H and the flexible 6-arm-linker were mixed into the solution first and allowed to bind for 10 min. As a last step before encapsulation, the G monomer was added.

### Preparation of 6-arm DNA-HMPs

Formation of the 6-arm DNA-HMPs was likewise achieved using droplet-templated formation in water-in-oil droplets using the monomers G and H and the 6-arm-linker in 6x excess. The droplets containing the DNA-solution were then placed into a thermal cycler (BioRad) and the DNA inside annealed by heating the samples to 78*^◦^*C for 15 min and subsequently cooling to 20*^◦^*C at an increment rate of 0.1*^◦^*C/min. Note that in order to prevent droplet fusion during heating in the thermal cycler, the surfactant perfluoropolyether–polyethylene glycol (PFPE–PEG, RAN Biotechnologies) was applied at 5 wt% instead of 2 wt% concentration.

### Confocal fluorescence microscopy

For imaging, an LSM 900 Zeiss confocal fluorescence microscope (Carl Zeiss AG) was used. For each experiment, the pinhole size was set to one Airy unit and either a Plan-Apochromat 20x/0.8 Air M27 objective or a 63x/1.2 W korr objective with oil immersion were utilized. All experiments were conducted at 22*^◦^*C room temperature. For imaging, the DNA-HMPs were deposited into custom-built observation chambers made from glass slides (Carl Roth) attached via double-sided sticky tape (Tesa) and sealed using two-component glue (twin-sil, Picodent). Prior to assembly of the observation chamber, the glass slides were coated for 5 min with poly(vinyl-alcohol) (50 mg*/*mL, Sigma Aldrich). For image analysis, ImageJ 1.54f (NIH, [62, 63]) was employed.

### Analysis of DNA-HMP size distributions

Analysis of DNA-HMP size distributions was conducted using the open-source software Fiji ImageJ [62, 63]. Confocal microscopy images were processed by first applying a Gaussian-filter to smooth particle outlines (*σ* = 2.0) followed by thresholding using Otsu’s method. The images were then binarized and a watershed algorithm applied to separate DNA-HMPs too close to one another. Particle size was then measured using “Analyze Particles” setting the size range to 50 µm^2^ to infinity and the circularity to 0 -1.0 saving Feret’s diameter as a size read-out. DNA-HMPs at the image-edge, which were only partially detected by this measurement, were discarded from the dataset.

### Fluorescence recovery after photobleaching

Fluorescence recovery after photobleaching (FRAP) experiments were conducted and analyzed as previously outlined [36].

### Analysis of DNA-HMP formation over time

The formation of DNA-HMPs (20 µM concentration, 3-arm short design) was tracked via confocal fluorescence microscopy over the course of 2 h using 5x digital zoom and 4x line averaging at an interval of 30 s. Per experiment, the formation of one DNA-HMP was tracked this way and the experiment conducted three times. The area fraction of the fluorescent signal inside the water-in-oil droplets was then analyzed using ImageJ 1.54f (NIH, [62, 63]). The images of the individual droplets were then binarised using Otsu’s method. Utilizing the circle tool to cover the whole droplet imaged, the area fraction of the fluorescence pixels over all pixels within the selected area was measured for each frame. The measured area fractions were normalized to the first frame of each video and the average of these values (n = 3 DNA-HMPs ± standard deviation) plotted over time.

Next, the formation of the DNA-HMPs following the condensation of DNA aggregates formed across the whole water-in-oil droplet was analyzed. For this, all frames were filtered using a Gaussian blur filter (*σ* = 2.0) to improve the signal to noise ratio. Using the plot profile function in ImageJ, whole droplet profiles were taken at nine different time points across the recorded experiment using the square tool to encase the whole droplet. Profiles were plotted in OriginPro 2021 - Update 6 (Origin Lab Corporation) and their peaks counted and analyzed using its Peak Analyzer tool to get the number of peaks and the gray value at each peak’s center for every recorded profile, which were then plotted over the recorded time.

### Cryogenic scanning electron microscopy (CryoSEM)

CryoSEM sample preparation was performed as previously described [64]. Briefly, 3 µL of a droplet emulsion was deposited onto a 0.8 mm-diameter gold specimen carrier mounted on a freeze-fracture holder (Leica Microsystems) and immediately immersed in liquid nitrogen. The cryo-embedded droplets were then transferred using an evacuated, liquid nitrogen-cooled shuttle (Leica EM VCT100, Leica Microsystems) into a freeze-fracture and etching system (Leica EM BAF060, Leica Microsystems). Fracturing was carried out in a vacuum chamber (10^-6^to 10^-7^mbar) at −160*^◦^*C using a cooled knife. To enable sublimation of water from the fractured droplets, the sample stage was subsequently heated to −90*^◦^*C for 45 min. After sublimation, the freeze-fractured droplets were coated with a 4 nm layer of platinum/carbon (Pt–C) by electron beam evaporation. For image acquisition, the samples were transferred via the same liquid nitrogen-cooled shuttle into the imaging chamber of a field-emission scanning electron microscope (FE-SEM; Zeiss Ultra 55, Carl Zeiss Microscopy), equipped with in-lens, secondary electron (SE), and angle-selective back-scattered electron (ASB) detectors (Carl Zeiss SMT). Top-view imaging was performed under cryogenic conditions (stage temperature −115*^◦^*C ± 5*^◦^*C) with a working distance of 3 - 5 mm. Due to the low conductivity of the emulsion droplets, low acceleration voltages of 1.5 – 4.0 kV were used. Signals were detected using the in-lens detector.

### Sample preparation and workflow for microindentation

24 Well Glass bottom Plates (Cellvis) were treated with oxygen plasma under vacuum (0.5 mbar final pressure, 200 W) for 3 min using a 300 Semi-auto plasma processor (PVA TePla AG). The wells were next filled with 1 mL of a 0.5 mg*/*mL poly-l-lysine (MW = 150 kDa - 300 kDa, Sigma Aldrich) solution in MilliQ water, and incubated for 60 min. The wells were then washed three times using 1x PBS and 10 mM MgCl_2_ solution. 100 µL of DNA-HMPs in solution were added to a total volume of 1 mL 1x PBS and 10 mM MgCl_2_ in the wells and allowed to settle and adhere to the poly-l-lysine coated wells for 30 min before washing the wells three times using 1x PBS and 10 mM MgCl_2_ solution. Indentation experiments were conducted on a Pavone microindenter (Optics11Life) using dynamic mechanical analysis (DMA) in displacement mode. For the single particle measurements, a cantilever with a tip size of 3.5 µm and a spring constant of 0.019 N*/*m was used. DMA was conducted at 0.1, 0.5, 1, 2, 5, 10 and 20 Hz frequency applying an indentation amplitude of 100 nm, 2 s relaxation time between frequencies and an initial relaxation time of 10 s. For each sample, measurements on five separate DNA-HMPs were undertaken. Each condition was measured as triplicates. Calculation of the Young’s moduli was based on Hertz-model fits of the indentation curve fitted to an indentation depth equal to 16% of the cantilever-tip diameter (560 nm). Further, each sample was indented no further than 5% of the sample diameter, to exclude any measurement artifacts based on the underlying substrate. The data is presented as the mean ± the standard deviation of the resulting values for the Young’s modulus. Storage, loss moduli and tan(*δ*) of the different types of DNA-HMPs are presented as single data points. Data analysis was conducted using the analysis software DataViewer (V2.5.0, Optics11Life), while plotting of the data as well as the statistical analysis were managed using OriginPro 2021 - Update 6 (Origin Lab Corporation).

### Measurement of hydrodynamic drag forces during microindentation

Oscillating cantilevers in viscous medium experience a force even in the absence of physical contact with a surface due to hydrodynamic drag [65, 66]. This drag depends on the viscosity and density of the medium, the size of the cantilever, and the frequency of oscillation. Thus, it needs to be determined for any given set of experimental parameters, as it can otherwise be mistakenly attributed to the properties of the material being tested during DMA. We thus measured the hydrodynamic drag force experienced by the utilized cantilever to estimate its influence on DMA measurements and corrected the data accordingly. For this, the cantilever tip was placed 500 nm above a DNA-HMP and the DMA sweep from 0.1 - 20 Hz performed there to estimate the viscous drag. This measurement was repeated five times to yield an average drag force measurement. For a detailed description on the correction of the drag force, see Supplementary Note 1 and Figures S20/S21.

### Sample preparation and workflow for real-time deformability cytometry

Real-time deformability cytometry (RT-DC) was performed using an AcCellerator (Zellmechanik Dresden) on an inverted AxioObserver microscope (Carl Zeiss AG) equipped with a 20x/0.4 Ph2 Plan-NeoFluar objective (Carl Zeiss AG). Images were acquired using a high-speed CMOS-camera (MC1362, Microtron).

Right before each measurement, DNA-HMPs in suspension (100 µL per run) were pelleted for 1 min using a C1008-GE myFUGE mini centrifuge (Benchmark Scientific). 80 µL of supernatant were discarded and the pellet resuspended in 150 µL CellCarrierB (Zellmechanik Dresden). The resuspended DNA-HMPs were then aspirated into a 1 mL glass syringe with PEEK tubing connector and PTFE plunger (SETonic) mounted on a syringe pump system (NemeSys, Cetoni). The DNA-HMP-CellCarrierB solution was then applied to a Flic20 microfluidic chip (Zellmechanik Dresden) using PTFE-tubing (S1810-12, Bola). Through a second 1 mL glass syringe, CellCarrierB was applied to the Flic20 microfluidic chip as sheath flow for the RT-DC experiment. For all samples, two sequential measurements using 0.04 µL*/*s and 0.08 µL*/*s total flow rates (ratio of sheath to sample flow: 3:1) were performed for a duration of at least 900 s each. The measurement software ShapeIn (version 2.2.2.4, Zellmechanik Dresden) was used to detect DNA-HMPs in real time. The pixel-size was adjusted to 0.68 µm/px, fitting the utilized 20x/0.4 Ph2 objective and all DNA-HMPs imaged at the rear part of the flow channel ensuring regular deformation of each particle. For each condition triplicates were measured. The analysis software Shape-Out (version 2.10.0, Zellmechanik Dresden) was then used to analyze the behavior of the DNA-HMPs in the flow channel. All samples were equally gated for porosity (1.0 - 1.15) and DNA-HMP size (40 µm^2^ - 180 µm^2^). Statistical significance was analyzed using a linear mixed model (R-lme4) as integrated in Shape-Out (version 2.10.0 [67]) yielding ANOVA p-values. Calculation of the Young’s modulus, deformation and volume as well as preparation of the data for contour plots were all carried out using Shape-Out (version 2.10.0). Population mean values of the measured samples are presented with their standard error of the mean. Plots for the volume, deformation and Young’s modulus were created using OriginPro 2021 - Update 6 (Origin Lab Corporation).

### Dynamic real-time deformability cytometry

For dynamic real-time deformability cytometry (dRT-DC), the width of the region in which images are taken during measurement was set to the maximum of 1200 px. Likewise, the frame rate of the CMOS-camera (MC1362, Microtron) was set to the maximum of 7000 f/s to allow for the imaging of individual particles as they undergo deformation. Sample preparation and data analysis was conducted in the same way as for the other RT-DC experiments. Plots and data fits were created using OriginPro 2021 - Update 6 (Origin Lab Corporation). To fit the deformation data of the DNA-HMPs during dRT-DC measurements, the speed of the particles inside of the flow channel was calculated. For this, the volume of the whole channel as 20 µm ∗ 20 µm ∗ 300 µm = 120 000 µm^3^ was taken into account. The volume of the channel divided by the flow rate of 0.04 µL*/*s as 40 000 000 µm^3^*/*s resulted in a channel traversing time of 3 ms for the DNA-HMPs. Given the channel length of 300 µm, the speed of the DNA-HMPs was calculated as 100 µm*/*ms and the data plotted accordingly. The relaxation (characteristic response) time *τ* as well as its standard error of the different DNA-HMP populations were then extracted from exponential fits of the resulting deformation curves using OriginPro 2021 - Update 6 (Origin Lab Corporation).

### Formation of c[RGD]-tagged DNA linker strands

The small peptide cyclo[Arg-Gly-Asp-D-Phe-Lys(Azide)] (c[RGD], Hölzel Diagnostika Handels GmbH) was diluted in a 1x PBS (no CaCl_2_, no MgCl_2_, pH = 7.4; Gibco) solution to yield a concentration of 2.4 mM. DBCO-tagged DNA strands (elongated linker DBCO, 6-arm-linker DBCO and flexible 6-arm-linker DBCO) were then dissolved at a final DNA concentration of 800 µM using the c[RGD]-solution creating a 3:1 ratio of azide to DBCO increasing the reaction yield. The solution was then incubated at 4*^◦^*C for 72 h to allow the DBCO-azide click reaction to occur. The resulting solution was then stored at −20*^◦^*C until further use.

### Polyacrylamide gel electrophoresis

Polyacrylamide gel electrophoresis (PAGE) was performed as previously described [36]. Here, however, a 15% polyacrylamide gel and a final concentration of 1x PBS (PBS, no CaCl_2_, no MgCl_2_, pH = 7.4; Gibco) was used as buffer in each sample. 10 µL Tridye Ultra low range DNA ladder (NEB) were used as reference. Gels were stained in 1× TBE (Tris Borate EDTA, Sigma, Thermo Scientific) buffer supplied with GelRed dye (Millipore) in a 1:10000 dilution for 10 min.

### Formation of c[RGD]-tagged DNA-HMPs

To create c[RGD]-tagged DNA-HMPs, 50 µL of the desired HMP suspension was pelleted for 1 min using a C1008-GE myFUGE mini centrifuge (Benchmark Scientific). Subsequently, 40 µL of the supernatant were removed and the DNA-HMP pellet resuspended using 10 µL of a 20 µM solution of the respective c[RGD]-tagged linker to yield a final concentration of 10 µM c[RGD]-tagged linker in the DNA-HMPs. HMPs were incubated with the linker overnight and subsequently washed 3x using a solution of 1x PBS and 10 mM MgCl_2_ to remove any non-incorporated peptides and DNA linkers.

### Incorporation of DNA-HMPs into 3D fibroblasts spheroids

DNA-HMPs were incorporated into 3D fibroblast spheroids, using the hanging drop technique, as described previously [60]. Briefly, primary mouse liver fibroblasts expressing td-Tomato (Mouse parental genotypes: TglnCreERT2/mTmG/TgfbrII lox/lox crossed with mTmG (Gt(ROSA)26Sortm4(ACTB-tdTomato,- EGFP)Luo/J), passage 20, mycoplasma negative) were used to form 3D cell spheroids. Every hanging drop comprised around 8500 fibroblasts and between 500 and 1000 DNA-HMPs. Cells and DNA-HMPs were co-cultured in cell media (DMEM, Gibco) supplemented with 1% FBS (Gibco), 1% penicillin-streptomycin (Gibco) and 10% methylcellulose (4000 cPs, Thermo Scientific) of 12 mg*/*mL concentration. Hanging drops of 15 µL volume were then formed at the inner part of the lid of a petri-dish, which was incubate at 37*^◦^*C under 5% CO_2_ atmosphere for 2 days or overnight. After that, the spheroids were collected and embedded in an alginate matrix of 2.5 mg*/*mL concentration, cross-linked with 0.12 M CaSO_4_ to keep their 3D and spherical shape during imaging. Primary mouse liver fibroblasts were acquired by the Trepat laboratory as a gift from the Batlle laboratory (IRB Barcelona). Originally, the cells were provided in passage 12.

### Imaging DNA-HMP deformation in 3D fibroblasts spheroids

DNA-HMP deformation inside 3D spheroids was tracked using a spinning disk confocal Dragonfly 200 (Andor) mounted on a Nikon Ti Eclipse microscope or a scanning confocal Zeiss LSM 900 microscope. Spheroids were captured using time-resolved z-stack imaging of several spheroids overnight, with frame rates of either 30, 45 or 60 min with a 100 µm total volume imaged per spheroid in steps of 1 µm, using a 20x Nikon objective. Image processing and video generation of the time-lapses were done using the open-source software ImageJ 1.54f (NIH, [62, 63]).

### Analysis of roundness and cross-section area of DNA-HMPs embedded in 3D fibroblast spheroids

Analysis of particle roundness and cross-section area of 4-arm, 6-arm flexible and 6-arm DNA-HMPs embedded in 3D fibroblast spheroids was conducted using ImageJ 1.54f (NIH, [62, 63]). Time-resolved z-stacks were loaded and substacks of individual DNA-HMPs extracted. Per substack, the equatorial plane of a DNA-HMP was extracted and the images compiled into a new stack containing the equatorial plane of the particle across time. To yield particle roundness and area, the stacks were filtered using a Gaussian-filter to smooth DNA-HMP outlines (*σ* = 1.0) and thresholded (Otsu’s method), yielding binarized images. Particles were then measured using “Analyze Particles”, setting a size range of 50 µm^2^ to infinity and a circularity range of 0.0 - 1.0, extracting the mean gray value, area and shape descriptors of the analyzed objects. To account for the low intensity of the 3-arm DNA-HMPs, corresponding data of this design was processed by removing noise via despeckling, smooth-filtering and thresholding (Otsu’s method) prior to particle analysis.

### Data analysis and statistics

All data apart from RT-DC data is presented showing the mean ± standard deviation of the given populations. RT-DC data is presented as the mean ± the standard error of the mean to account for the large sample sizes. In addition, the sample size is given for all depicted data. If not stated otherwise, statistical analysis was conducted using unpaired, two-tailed Student’s t-tests and a minimum p-value of 0.05 used to determine the statistical significance of the data. OriginPro 2021 - Update 6 (Origin Lab Corporation) was used to conduct the statistical tests and to prepare the presented graphs for all data other than RT-DC data. For RT-DC data, statistical significance was analyzed using a linear mixed model (R-lme4) as integrated in Shape-Out (version 2.10.0 [67]) yielding ANOVA p-values.

## Acknowledgements

K.G. acknowledges funding from the Deutsche Forschungsgemeinschaft (DFG, German Research Foundation) under Germany’s Excellence Strategy via the Excellence Cluster 3D Matter Made to Order (EXC-2082/1 – 390761711), the Human Frontiers Science Program (RGPO03I2023), the Alfried Krupp Förderpreis and the ERC Starting Grant “ENSYNC” (No. 101076997). T.W. thanks the Studienstiftung des deutschen Volkes e.V. The authors thank Dr. Sadaf Pashapour and the Microfabrication and Microfluidic Core Facility (µFluCF) at the Institute of Molecular Systems Engineering and Advanced Materials (IMSEAM) funded partly by the Health + Life Science Alliance Heidelberg Mannheim. The Health + Life Science Alliance provided state funds approved by the State Parliament of Baden-Württemberg. This paper was additionally funded by the Generalitat de Catalunya (AGAUR SGR-2017-01602 to X.T., the CERCA Programme, and “ICREA Academia” award to P.R-C.); the Spanish Ministry for Science and Innovation MICCINN/FEDER (PID2021-128635NB-I00, MCIN/AEI/ 10.13039/501100011033 and “ERDF-EU A way of making Europe” to X.T., PID2019-110298GB-I00 to P.R-C.); the European Research Council (Adv-101097753 to P.R-C. and Adv-883739 to X.T.); La Caixa Foundation (LCF/PR/HR24/00326 to X.T.); and the Human Frontiers Science Program (HFSPRGP022/2024). E.D. received funding from the Instituto de Salud Carlos III (Reference: IHMC22/00021). G.F. acknowledges funding from the Marie Sklodowska Curie Action (grant agreement 101206469). We would like to thank the Soft (bio)materials characterization Core Facility (Biomechanics) at IMSEAM Heidelberg University for allowing us to use their Pavone microindenter (Optics11Life), funded by the Federal Ministry of Education and Research (BMBF) and the Ministry of Science Baden-Württemberg within the framework of the Excellence Strategy of the Federal and State Governments of Germany. In particular, the authors thank Prof. Dr. Christine Selhuber-Unkel and Florine Sessler for access to and training on the Pavone microindenter. The authors thank our technician Claudia Helbig for assistance with PAGE. The authors thank Tobias Abele for continued discussions throughout the duration of the project. The authors thank Dr. Bob Fregin and Prof. Dr. Oliver Otto, University of Greifswald, for discussions on the analysis of the dRT-DC data.

