## Supplementary Information for "Mechanically Tunable DNA Hydrogel Microparticles for 3D Cellular Systems"

Dr. Gotthold Fläschner, Prof. Dr. Pere Roca-Cusachs
Address: Institute for Bioengineering of Catalonia (IBEC), Cellular and molecular mechanobiology, Baldiri Reixac 15-21, 08028 Barcelona

Dr. Sadaf Pashapour
Address: Heidelberg University, Institute for Molecular Systems Engineering and Advanced Materials (IMSEAM), Heidelberg University
Microfabrication and Microfluidics Core Facility (µFluCF), Institute for Molecular Systems Engineering and Advanced Materials (IMSEAM), Heidelberg University, INF 225, 69120 Heidelberg

Dr. Ilia Platzman
Address: Max Planck Institute for Medical Research, Department of Cellular Biophysics, Jahnstraße 29, 69120 Heidelberg, Germany

4 Supplementary Note 1: Correction for drag force 28

### 1 Supporting Tables

#### 1.1 Table S1: List of DNA sequences

Table S1: DNA sequences used in this study. The fluorescent label cyanine 3 is abbreviated as Cy3. Dibenzocyclooctin is abbreviated as DBCO. Modifications are highlighted in italic.

| **Name** | **DNA sequence 5' - 3'** |
|---|---|
| **3-arm short** | |
| A-1 | GACCAACACCAGTGAGGACGGAAGTTTGTCGTAGCATCGCACC |
| A-2 | GACCAACACCAACCACGCCTGTCCATTACTTCCGTCCTCACTG |
| A-3 | GACCAACACGGTGCGATGCTACGACTTTGGACAGGCGTGGTTG |
| B-1 | CAGTGAGGACGGAAGTTTGTCGTAGCATCGCACCCGACAGGAA |
| B-1-Cy3 | *Cy3*-CAGTGAGGACGGAAGTTTGTCGTAGCATCGCACCCGACAGGAA |
| B-2 | CAACCACGCCTGTCCATTACTTCCGTCCTCACTGCGACAGGAA |
| B-3 | GGTGCGATGCTACGACTTTGGACAGGCGTGGTTGCGACAGGAA |
| Linker | GTGTTGGTCTTCCTGTCG |
| **3-arm** | |
| C-1 | TGCGACCAACACCAGTGAGGACGGAAGTTTGTCGTAGCATCGCACC |
| C-2 | TGCGACCAACACCAACCACGCCTGTCCATTACTTCCGTCCTCACTG |
| C-3 | TGCGACCAACACGGTGCGATGCTACGACTTTGGACAGGCGTGGTTG |
| D-1 | CAGTGAGGACGGAAGTTTGTCGTAGCATCGCACCACGCGACAGGAA |
| D-1-Cy3 | *Cy3*-CAGTGAGGACGGAAGTTTGTCGTAGCATCGCACCACGCGACAGGAA |
| D-1-390 | *Atto390*-CAGTGAGGACGGAAGTTTGTCGTAGCATCGCACCACGCGACAGGAA |
| D-2 | CAACCACGCCTGTCCATTACTTCCGTCCTCACTGACGCGACAGGAA |
| D-3 | GGTGCGATGCTACGACTTTGGACAGGCGTGGTTGACGCGACAGGAA |
| Elongated Linker | GTGTTGGTCGCATTCCTGTCGCGT |
| Elongated Linker DBCO | *DBCO*-TGTGTTGGTCGCATTCCTGTCGCGT |
| **4-arm** | |
| E-1 | CTACTATGGCGGGTGATAAATTCGGGAAGAGCATGCCCATCCACGCGACAGGAA |
| E-1-488 | *Atto488*-CTACTATGGCGGGTGATAAATTCGGGAAGAGCATGCCCATCCACGCGACAGGAA |
| E-2 | GGATGGGCATGCTCTTCCCGTTCTCAACTGCCTGGTGATACGACGCGACAGGAA |
| E-3 | CGTATCACCAGGCAGTTGAGTTCATGCGAGGGTCCAATACCGACGCGACAGGAA |
| E-4 | CGGTATTGGACCCTCGCATGTTTTTATCACCCGCCATAGTAGACGCGACAGGAA |
| F-1 | TGCGACCAACACCTACTATGGCGGGTGATAAATTCGGGAAGAGCATGCCCATCC |
| F-2 | TGCGACCAACACGGATGGGCATGCTCTTCCCGTTCTCAACTGCCTGGTGATACG |
| F-3 | TGCGACCAACACCGTATCACCAGGCAGTTGAGTTCATGCGAGGGTCCAATACCG |
| F-4 | TGCGACCAACACCGGTATTGGACCCTCGCATGTTTTTATCACCCGCCATAGTAG |
| **6-arm** | |
| G-1 | GCTGGACTAACGGAACGGTTAGTCAGGTATGCCAGCACATAGCTTCCTCG |
| G-1-647 | *Atto647N*-GCTGGACTAACGGAACGGTTAGTCAGGTATGCCAGCACATAGCTTCCTCG |
| G-2 | CTCAGAGAGGTGACAGCATTCCGTTCCGTTAGTCCAGCATAGCTTCCTCG |
| G-3 | CCATGGTCCCAAGTGATGTTTGCTGTCACCTCTCTGAGATAGCTTCCTCG |
| G-4 | CGGCGCTGTAAATTTGCGTTCATCACTTGGGACCATGGATAGCTTCCTCG |
| G-5 | CAGACGTCACTCTCCAACTTCGCAAATTTACAGCGCCGATAGCTTCCTCG |
| G-6 | GTGCTGGCATACCTGACTTTGTTGGAGAGTGACGTCTGATAGCTTCCTCG |
| H-1 | TGCGACCAACACGCTGGACTAACGGAACGGTTAGTCAGGTATGCCAGCAC |
| H-2 | TGCGACCAACACCTCAGAGAGGTGACAGCATTCCGTTCCGTTAGTCCAGC |
| H-3 | TGCGACCAACACCCATGGTCCCAAGTGATGTTTGCTGTCACCTCTCTGAG |
| H-4 | TGCGACCAACACCGGCGCTGTAAATTTGCGTTCATCACTTGGGACCATGG |
| H-5 | TGCGACCAACACCAGACGTCACTCTCCAACTTCGCAAATTTACAGCGCCG |
| H-6 | TGCGACCAACACGTGCTGGCATACCTGACTTTGTTGGAGAGTGACGTCTG |
| 6-arm-linker | GTGTTGGTCGCACGAGGAAGCTAT |
| Flexible 6-arm-linker | GTGTTGGTCGCATTCGAGGAAGCTAT |
| 6-arm-linker DBCO | *DBCO*-TGTGTTGGTCGCACGAGGAAGCTAT |
| Flexible 6-arm-linker DBCO | *DBCO*-TGTGTTGGTCGCATTCGAGGAAGCTAT |

#### 2 Supporting Figures

##### 2.1 Figure S1: Designs of microfluidic devices used to create size-controlled DNA-HMPs

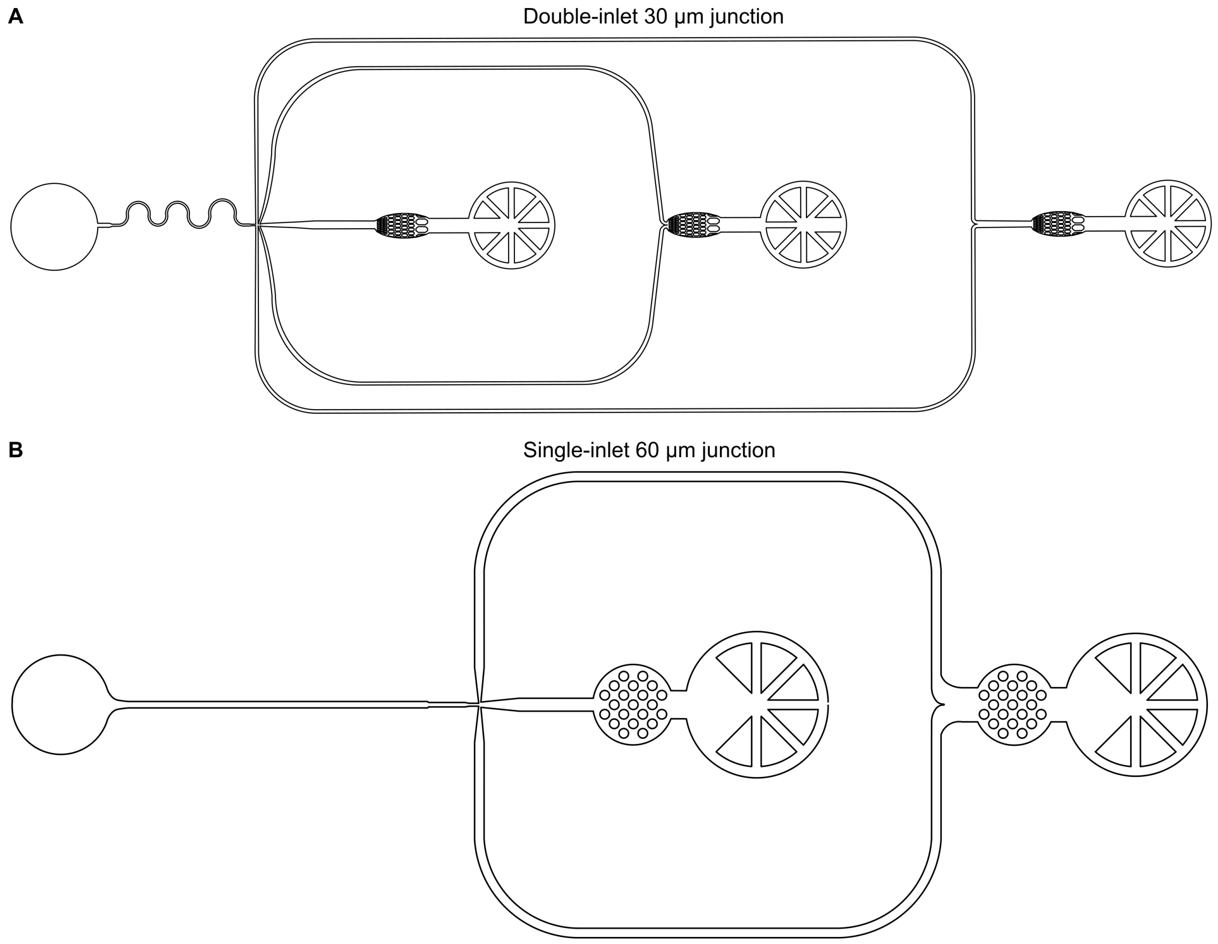

Figure S1: Microfluidics chips used for the preparation of size-controlled DNA-HMPs. A) 30 µm junction double-inlet device used to create DNA-HMPs of sizes up to 30 µm. B) 60 µm junction single-inlet device used to create DNA-HMPs of sizes up to 60 µm.

#### 2.2 Figure S2: Time- and linker dependent formation of DNA-HMPs

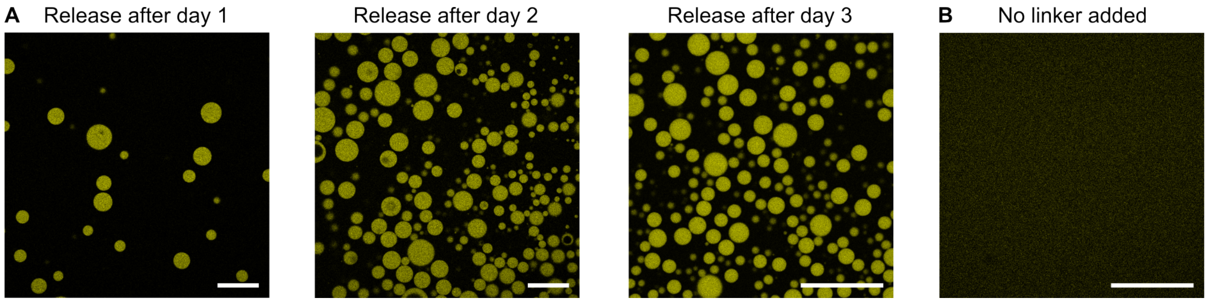

Figure S2: Control experiments of DNA-HMP formation. A) Confocal microscopy ($\lambda_{ex} = 561\,\text{nm}$, Cy3-labeled DNA, yellow) images of 3-arm short DNA-HMPs released after 1, 2 and 3 days of incubation in water-in-oil droplets. The highest yield of intact DNA-HMPs was achieved after 3 day incubation. B) No intact DNA-HMPs can be released from water-in-oil droplets following incubation in water-in-oil droplets of the nanostars without the linker strand. Scale bars: 100 µm.

#### 2.3 Figure S3: Droplet-templated formation of 3-arm short DNA-HMPs over time

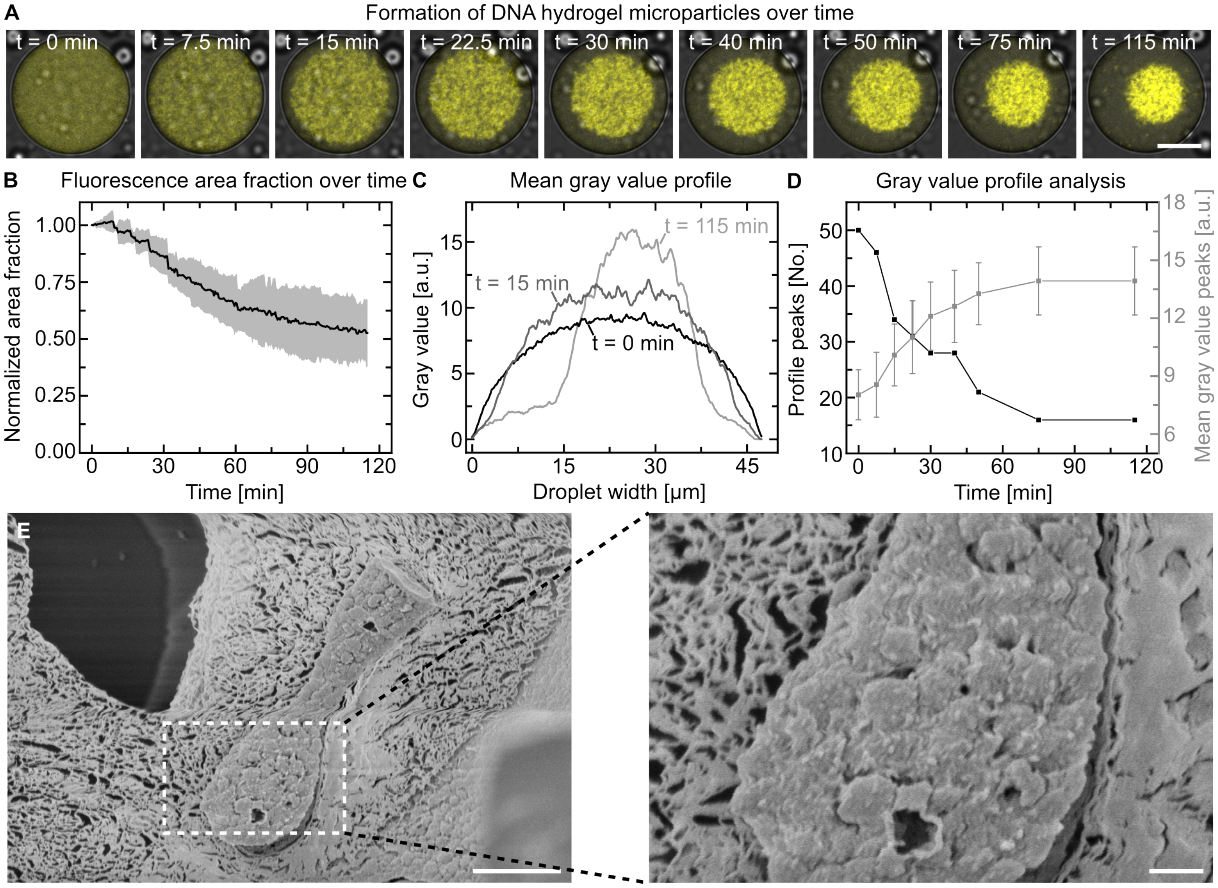

Figure S3: Formation of DNA-HMPs inside of water-in-oil droplets over time. A) Overlay of confocal microscopy ($\lambda_{ex} = 561$ nm, Cy3-labeled DNA, yellow) and brightfield images of a 3-arm short DNA-HMP forming inside a water-in-oil droplet over the course of 115 min, showing how the DNA condensates simultaneously across the whole volume of the water-in-oil droplet to form a singular DNA-HMP. Scale bar: 20 µm. B) Measured fluorescence area fraction of the DNA signal inside of water-in-oil droplets over time. The mean measured area fraction of fluorescent pixels over all pixels within the acquired droplet images is shown, depicting a decrease in area until the resulting DNA-HMP is formed (n = 3, ± standard deviation). C) Exemplary gray value profiles from three time points of the images shown in A, showing an increase in fluorescence intensity over time as well as a narrowing of the distribution indicating concentration of signal. D) Analysis of all gray value profiles of the images shown in A. Over time, the number of fluorescence peaks measured across the droplet decreases (black curve, total number of detected curve peaks at each time point), while the mean gray value of the fluorescence peaks increases (light gray curve, mean ± standard deviation of each measured time point). Both plateau towards the end of the particle formation, depicting the condensation of the DNA across the droplet towards its center over time. E) Cross-section of a 3-arm short DNA-HMP at 20 µM DNA concentration as seen via cryoSEM imaging. In the center and on the right-hand side of the image, uncut regions of the network are visible showing phase-separated droplets of DNA of several nanometers in size. Scale bar: 1 µm. The zoom shows the phase-separated region in the center of the droplet in more detail. Scale bar: 200 nm.

#### 2.4 Figure S4: Confocal microscopy of DNA-HMPs created using 3-arm, 4-arm, 6-arm and 6-arm flexible DNA nanostars

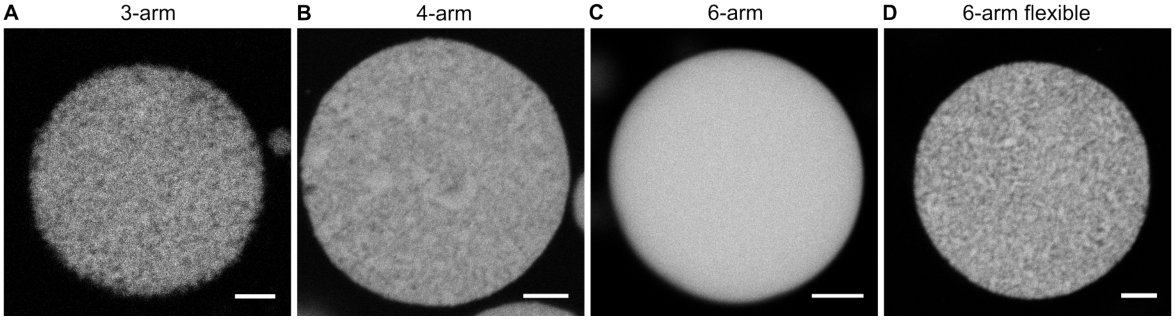

Figure S4: Microscopy data showing DNA-HMPs created using 3-arm, 4-arm, 6-arm and 6-arm flexible nanostars. A) Confocal microscopy ($\lambda_{ex} = 561\,\text{nm}$, Cy3-labeled DNA) image of a 3-arm DNA-HMP. B) Confocal microscopy ($\lambda_{ex} = 488\,\text{nm}$, Atto488-labeled DNA) image of a 4-arm DNA-HMP. C) Confocal microscopy ($\lambda_{ex} = 640\,\text{nm}$, Atto647N-labeled DNA) image of a 6-arm DNA-HMP. D) Confocal microscopy ($\lambda_{ex} = 640\,\text{nm}$, Atto647N-labeled DNA) image of a 6-arm flexible DNA-HMP. Scale bars: 10 µm.

#### 2.5 Figure S5: Binding of DNA-HMP to a glass substrate after poly-l-lysine functionalization following electrostatic interaction

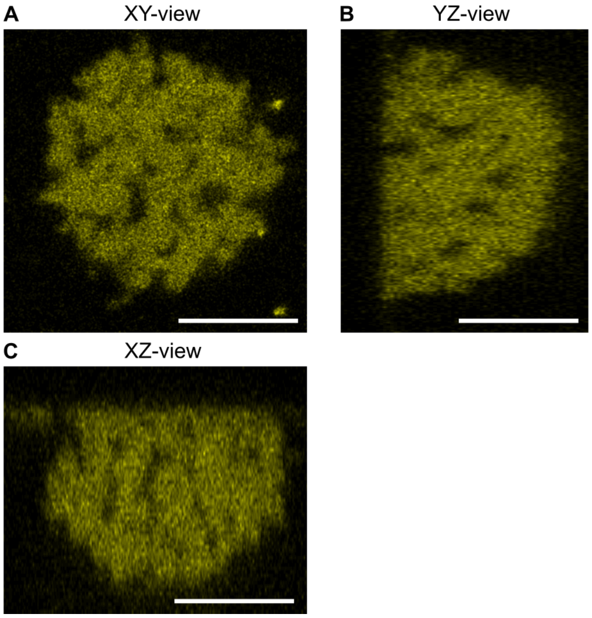

Figure S5: DNA-HMP bound to glass substrate following poly-l-lysine functionalization. A) Confocal microscopy ($\lambda_{ex} = 561$ nm, Cy3-labeled DNA, yellow) image of the xy-view of a 3-arm short DNA-HMP z-stack following electrostatic binding of the particle to a poly-l-lysine functionalized glass surface. B) Confocal microscopy ($\lambda_{ex} = 561$ nm, Cy3-labeled DNA, yellow) image of the yz-view of a 3-arm short DNA-HMP z-stack following electrostatic binding of the particle to a poly-l-lysine functionalized glass surface. C) Confocal microscopy ($\lambda_{ex} = 561$ nm, Cy3-labeled DNA, yellow) image of the xz-view of a 3-arm short DNA-HMP z-stack following electrostatic binding of the particle to a poly-l-lysine functionalized glass surface. Scale bars: 10 µm.

#### 2.6 Figure S6: Dynamic mechanical analysis of 3-arm short DNA-HMPs

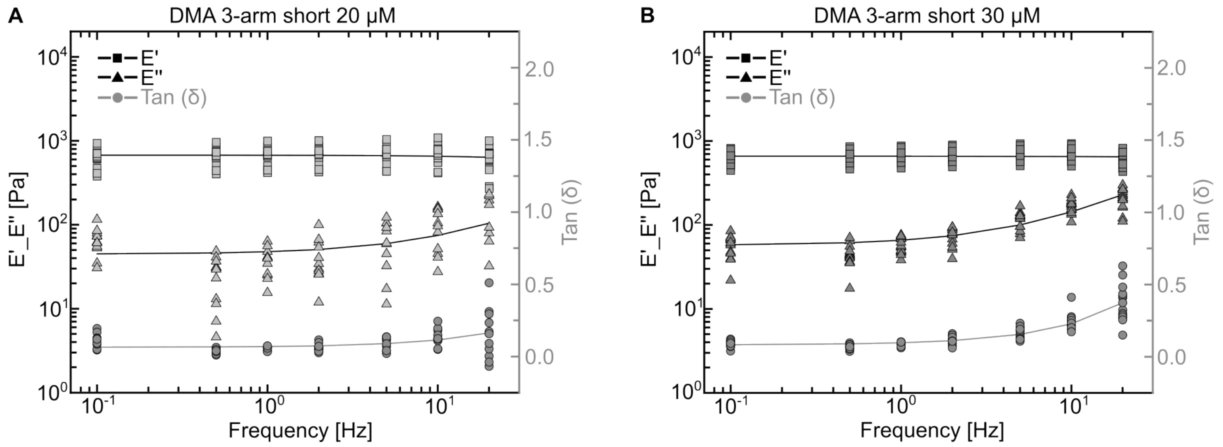

Figure S6: Further dynamic mechanical analysis (DMA) of 3-arm short DNA-HMPs. A) DMA plot of 3-arm short DNA-HMPs at 20 µM DNA concentration. The storage modulus E' (squares) and the loss modulus E" (triangles) are plotted on a logarithmic scale, tan ($\delta$) (E"/E', filled circles) is plotted on a linear scale. Tan ($\delta$) stays almost constant across all measured frequencies and only increases slightly at the highest measured frequency denoting the 3-arm short DNA-HMPs as behaving predominantly elastic. B) DMA plot of 3-arm short DNA-HMPs at 30 µM DNA concentration. The storage modulus E' (squares) and the loss modulus E" (triangles) are plotted on a logarithmic scale, tan ($\delta$) (E"/E', filled circles) is plotted on a linear scale.

#### 2.7 Figure S7: Dynamic mechanical analysis of 3-arm DNA-HMPs

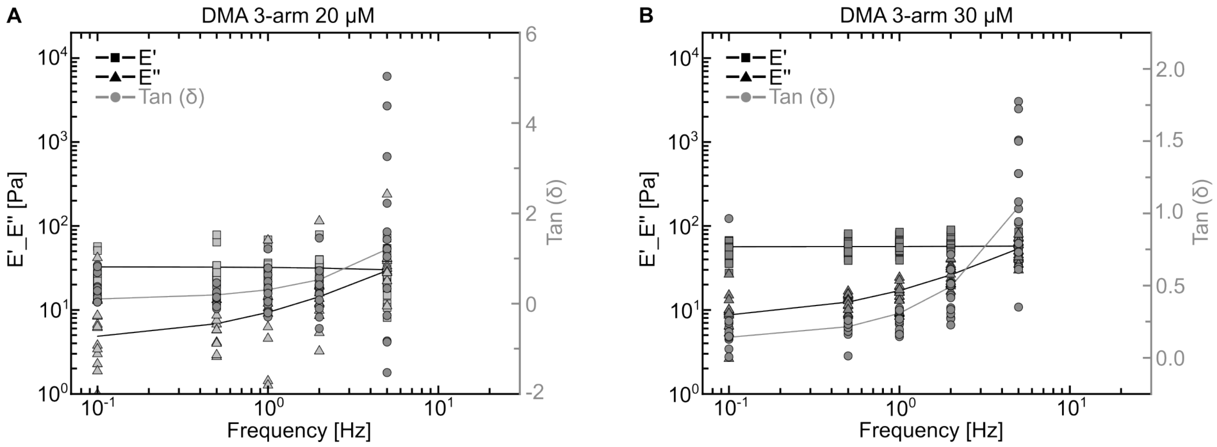

Figure S7: Further dynamic mechanical analysis (DMA) of 3-arm DNA-HMPs. A) DMA plot of 3-arm DNA-HMPs at 20 µM DNA concentration. The storage modulus E' (squares) and the loss modulus E" (triangles) are plotted on a logarithmic scale, tan ($\delta$) (E"/E', filled circles) is plotted on a linear scale. Tan ($\delta$) can be seen to increase more strongly already at lower frequencies showing clearly the cross-over point of E' and E" at 5 Hz, denoting the 3-arm DNA-HMPs as showing a more viscous response to higher mechanical stress than the 3-arm short DNA-HMPs. B) DMA plot of 3-arm DNA-HMPs at 30 µM DNA concentration. The storage modulus E' (squares) and the loss modulus E" (triangles) are plotted on a logarithmic scale, tan ($\delta$) (E"/E', filled circles) is plotted on a linear scale.

#### 2.8 Figure S8: Dynamic mechanical analysis of 4-arm DNA-HMPs

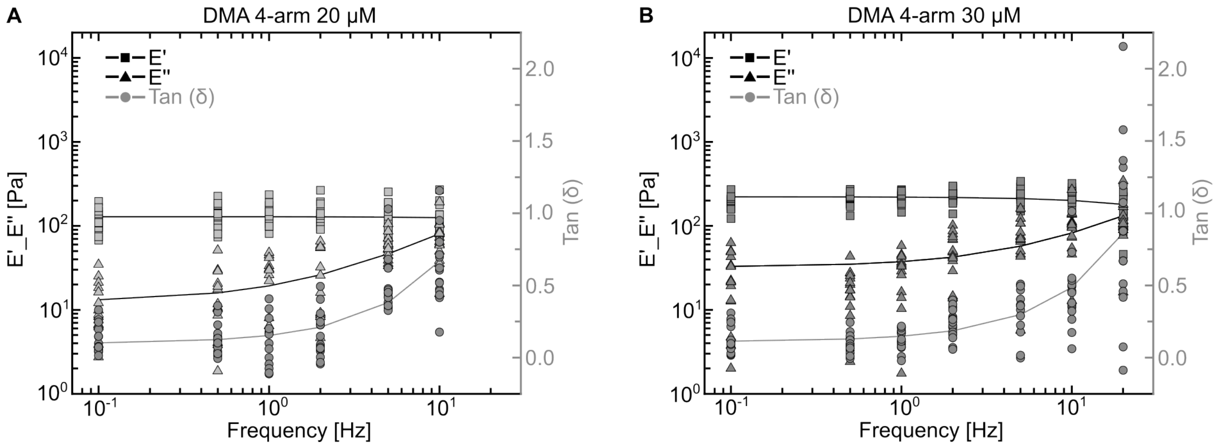

Figure S8: Further dynamic mechanical analysis (DMA) of 4-arm DNA-HMPs. A) DMA plot of 4-arm DNA-HMPs at 20 µM DNA concentration. The storage modulus E' (squares) and the loss modulus E" (triangles) are plotted on a logarithmic scale, tan ($\delta$) (E"/E', filled circles) is plotted on a linear scale. Tan ($\delta$) can be seen to increase more strongly already at lower frequencies showing clearly the cross-over point of E' and E" at 10 Hz, denoting the 4-arm DNA-HMPs as showing a more viscous response to higher mechanical stress than the 3-arm short DNA-HMPs, but less viscous than the 3-arm DNA-HMPs. B) DMA plot of 4-arm DNA-HMPs at 30 µM DNA concentration. The storage modulus E' (squares) and the loss modulus E" (triangles) are plotted on a logarithmic scale, tan ($\delta$) (E"/E', filled circles) is plotted on a linear scale.

#### 2.9 Figure S9: Dynamic mechanical analysis of 6-arm flexible DNA-HMPs

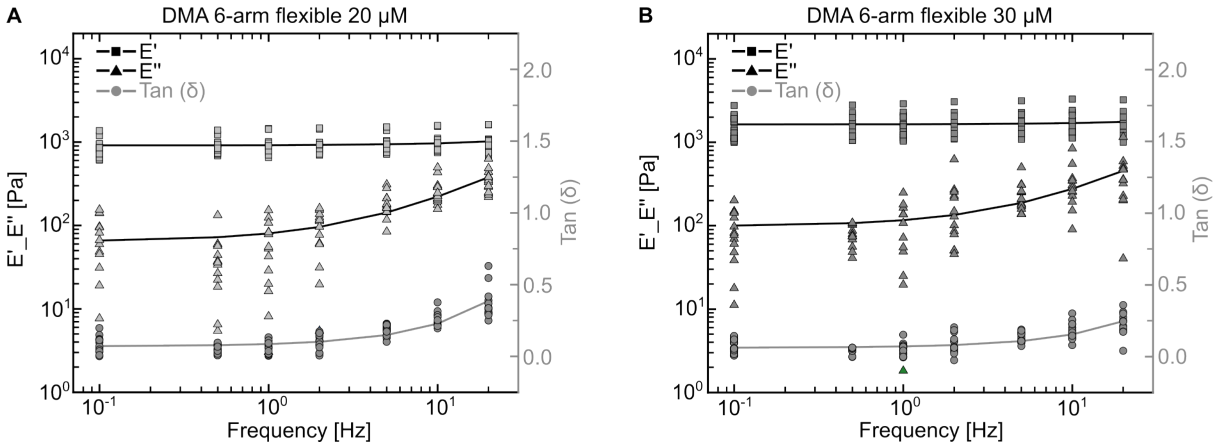

Figure S9: Further dynamic mechanical analysis (DMA) of 6-arm flexible DNA-HMPs. A) DMA plot of 6-arm flexible DNA-HMPs at 20 µM DNA concentration. The storage modulus E' (squares) and the loss modulus E" (triangles) are plotted on a logarithmic scale, tan ($\delta$) (E"/E', filled circles) is plotted on a linear scale. Tan ($\delta$) stays below 0.5 for all measured conditions with only a slight increase of E" at higher frequencies, denoting the 6-arm flexible DNA-HMPs as behaving mostly elastic. B) DMA plot of 6-arm flexible DNA-HMPs at 30 µM DNA concentration. The storage modulus E' (squares) and the loss modulus E" (triangles) are plotted on a logarithmic scale, tan ($\delta$) (E"/E', filled circles) is plotted on a linear scale.

#### 2.10 Figure S10: Dynamic mechanical analysis of 6-arm DNA-HMPs

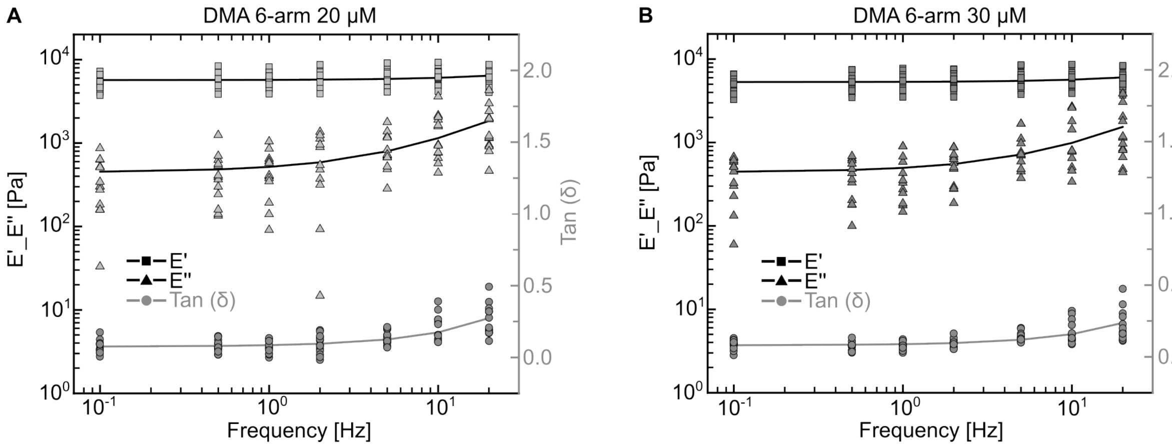

Figure S10: Further dynamic mechanical analysis (DMA) of 6-arm DNA-HMPs. A) DMA plot of 6-arm DNA-HMPs at 20 µM DNA concentration. The storage modulus E' (squares) and the loss modulus E" (triangles) are plotted on a logarithmic scale, tan ($\delta$) (E"/E', filled circles) is plotted on a linear scale. Tan ($\delta$) stays below 0.5 for all measured conditions with only a slight increase of E" at higher frequencies, denoting the 6-arm DNA-HMPs as behaving mostly elastic. B) DMA plot of 6-arm DNA-HMPs at 30 µM DNA concentration. The storage modulus E' (squares) and the loss modulus E" (triangles) are plotted on a logarithmic scale, tan ($\delta$) (E"/E', filled circles) is plotted on a linear scale.

#### 2.11 Figure S11: DNA-HMPs display different relaxation behaviors during microindentation

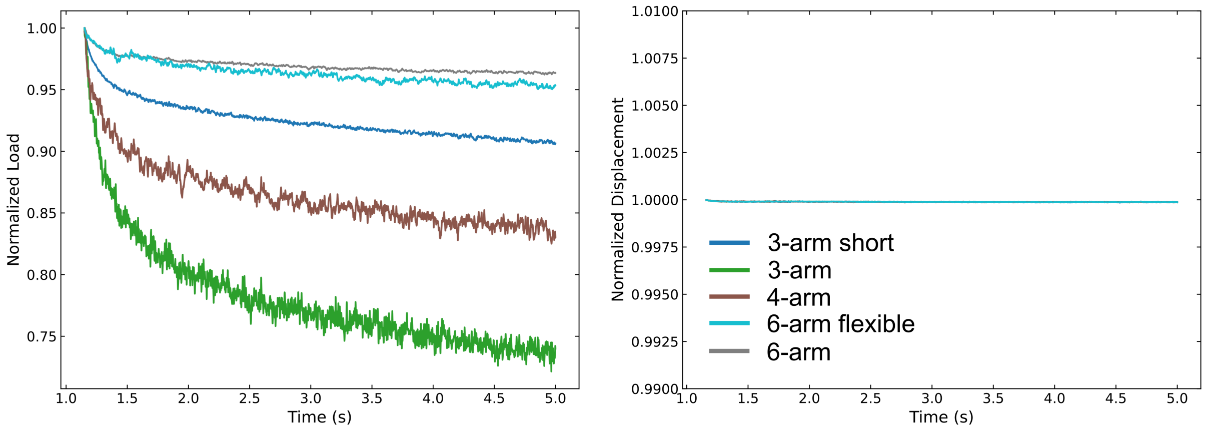

Figure S11: Relaxation behavior of DNA-HMPs across DNA nanostar designs. Directly after indentation prior to dynamic mechanical analysis, the DNA-HMPs are allowed to relax for 10 s. The normalized force detected by the cantilever during this relaxation period is plotted as a function of time for different DNA-HMP nanostars. 3-arm and 4-arm DNA-HMPs display a behavior consistent with a higher viscosity compared to the other designs (plot 1) with the 6-arm DNA-HMPs exhibiting the strongest elastic response. The normalized displacement of the cantilever is shown as a function of time for the same designs. Minimal initial displacement and variation are observed for all designs (plot 2).

#### 2.12 Figure S12: Deformation of DNA-HMPs during RT-DC

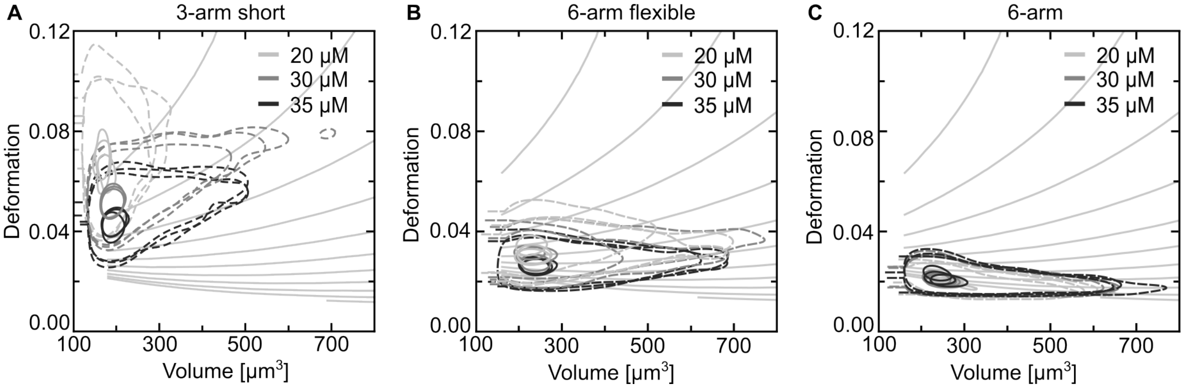

Figure S12: Full contour plots of DNA-HMP deformation during RT-DC showing all repeats. A) 3-arm short DNA-HMP deformation plotted against the volume for different DNA concentrations ($n_{20\,\mu M}$ = 45744, $n_{30\,\mu M}$ = 54771, $n_{35\,\mu M}$ = 52743). The data is presented using contour plots showing the $50^{th}$ percentile (dashed line) and $95^{th}$ percentile (solid line) of three independent measurements per condition. B) 6-arm flexible DNA-HMP deformation plotted against the volume for different DNA concentrations ($n_{20\,\mu M}$ = 23801, $n_{30\,\mu M}$ = 22632, $n_{35\,\mu M}$ = 18931). The data is presented using contour plots showing the $50^{th}$ percentile (dashed line) and $95^{th}$ percentile (solid line) of three independent measurements per condition. C) 6-arm DNA-HMP deformation plotted against the volume for different DNA concentrations ($n_{20\,\mu M}$ = 42430, $n_{30\,\mu M}$ = 42271, $n_{35\,\mu M}$ = 47844). The data is presented using contour plots showing the $50^{th}$ percentile (dashed line) and $95^{th}$ percentile (solid line) of three independent measurements per condition. Isoelasticity lines derived from numerical simulations are shown additionally, indicating stiffness changes where a steeper slope corresponds to softer particles.

#### 2.13 Figure S13: Real-time deformability cytometry of 3-arm short DNA-HMPs

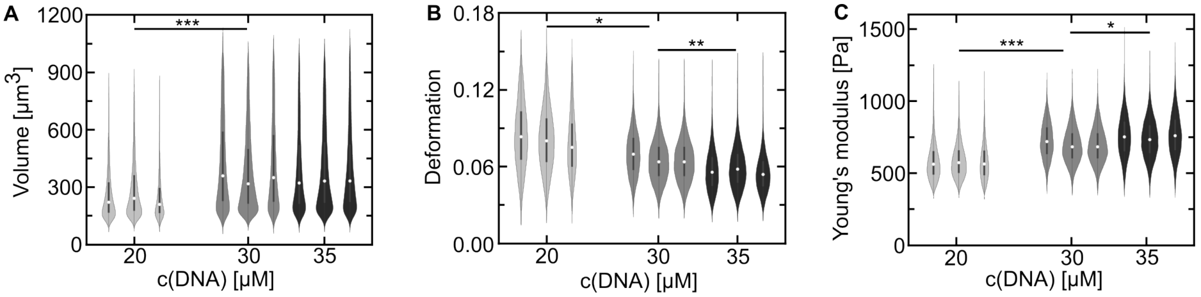

Figure S13: Analysis of 3-arm short DNA-HMPs using real-time deformability cytometry (RT-DC). A, B, C) Plots depicting changes in volume, deformation and Young's modulus of 3-arm short DNA-HMPs at 20 µM (n = 45744), 30 µM (n = 54771) and 35 µM (n = 52743) DNA concentration. The data is presented as violin plots showing the median value (white dot) of each measurement with boxplots encompassing the 25 - 75 % percentiles and a whisker length of 1.5 IQR. For each condition n = 3 independent experiments are displayed. DNA-HMP size increased significantly with an increase in DNA concentration from 20 µM, to 30 µM, while the size did not seem to change considerably when further increasing the DNA concentration to 35 µM (whole-population mean: $V_{20\,\mu M} = 273.7\,\mu m^3 \pm 2.1\,\mu m^3$, $V_{30\,\mu M} = 409.5\,\mu m^3 \pm 3\,\mu m^3$, $V_{35\,\mu M} = 397.1\,\mu m^3 \pm 2.9\,\mu m^3$). ***p-value: 0.0009. A significant decrease in deformation is shown for both 20 µM and 30 µM (also between 20 µM and 35 µM DNA-HMPs) as well as 30 µM and 35 µM DNA-HMPs (whole-population mean: $D_{20\,\mu M} = 0.081 \pm 0.0003$, $D_{30\,\mu M} = 0.069 \pm 0.0002$, $D_{35\,\mu M} = 0.058 \pm 0.0002$). *p-value: 0.015, ***p-value: 0.0005. Accordingly, the Young's modulus increased significantly between 20 µM and 30 µM DNA-HMPs as well as 30 µM and 35 µM DNA-HMPs (Whole population mean: $YM_{20\,\mu M} = 0.59\,kPa \pm 0.002\,kPa$, $YM_{30\,\mu M} = 0.71\,kPa \pm 0.002\,kPa$, $YM_{35\,\mu M} = 0.76\,kPa \pm 0.002\,kPa$). *p-value: 0.04. ***p-value: 0.0002. Statistical significance was analyzed using a linear mixed model (R-lme4) as integrated in Shape-Out (version 2.10.0) yielding ANOVA p-values. Error values correspond to the standard error of the mean.

#### 2.14 Figure S14: Real-time deformability cytometry of 6-arm flexible DNA-HMPs

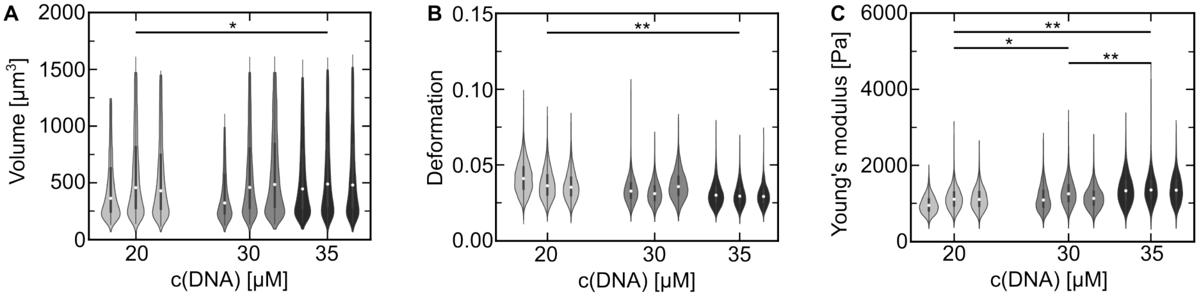

Figure S14: Analysis of 6-arm flexible DNA-HMPs using real-time deformability cytometry (RT-DC). A, B, C) Plots depicting changes in volume, deformation and Young's modulus of 6-arm flexible DNA-HMPs at 20 µM (n = 23801), 30 µM (n = 22632) and 35 µM (n = 18931) DNA concentration. The data is presented as violin plots showing the median value (white dot) of each measurement with boxplots encompassing the 25 - 75 % percentiles and a whisker length of 1.5 IQR. For each condition n = 3 independent experiments are displayed. DNA-HMP size increased significantly with an increase in DNA concentration from 20 µM to 35 µM (whole-population mean: $V_{20\,\mu M} = 540.4\,\mu m^3 \pm 6.3\,\mu m^3$, $V_{30\,\mu M} = 547.7\,\mu m^3 \pm 6.5\,\mu m^3$, $V_{35\,\mu M} = 591.9\,\mu m^3 \pm 6.8\,\mu m^3$). *p-value: 0.03. The deformation of the DNA-HMPs decreased significantly with an increase in DNA concentration from 20 µM to 35 µM (whole-population mean: $D_{20\,\mu M} = 0.039 \pm 0.0002$, $D_{30\,\mu M} = 0.035 \pm 0.0002$, $D_{35\,\mu M} = 0.031 \pm 0.0002$). **p-value: 0.0024. Likewise, the Young's modulus increased significantly from 20 µM to 35 µM between all concentrations (whole population mean: $YM_{20\,\mu M} = 1.09\,kPa \pm 0.006\,kPa$, $YM_{30\,\mu M} = 1.22\,kPa \pm 0.008\,kPa$, $YM_{35\,\mu M} = 1.41\,kPa \pm 0.009\,kPa$. *p-value: 0.034, **p-values: 0.007, 0.003. Statistical significance was analyzed using a linear mixed model (R-lme4) as integrated in Shape-Out (version 2.10.0) yielding ANOVA p-values. Error values correspond to the standard error of the mean.

#### 2.15 Figure S15: Real-time deformability cytometry of 6-arm DNA-HMPs

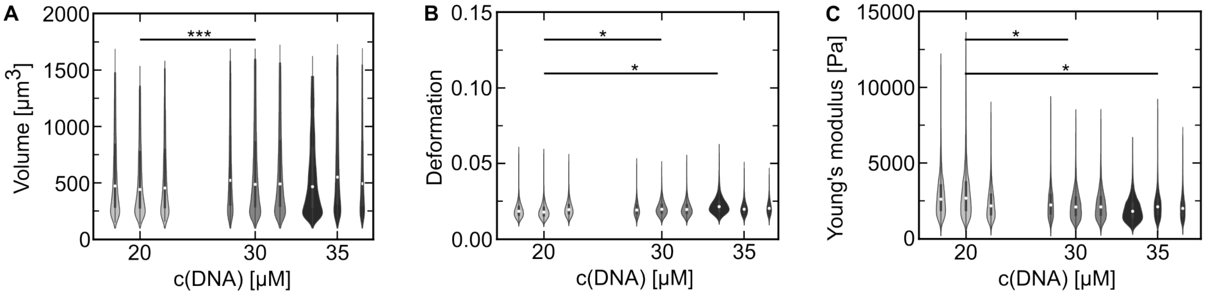

Figure S15: Analysis of 6-arm DNA-HMPs using real-time deformability cytometry (RT-DC). A, B, C) Plots depicting changes in volume, deformation and Young's modulus of 6-arm DNA-HMPs at 20 µM (n = 42430), 30 µM (n = 42271) and 35 µM (n = 47844) DNA concentration. The data is presented as violin plots showing the median value (white dot) of each measurement with boxplots encompassing the 25 - 75 % percentiles and a whisker length of 1.5 IQR. For each condition n = 3 independent experiments are displayed. DNA-HMP size increased significantly with an increase in DNA concentration from 20 µM to 30 µM (Whole-population mean volume: $V_{20\,\mu M} = 625.7\,\mu m^3 \pm 6.8\,\mu m^3$, $V_{30\,\mu M} = 629.1\,\mu m^3 \pm 7\,\mu m^3$, $V_{35\,\mu M} = 590.2\,\mu m^3 \pm 6.7\,\mu m^3$). ***p-value: 0.0002. The deformation of the DNA-HMPs increased significantly with an increase in DNA concentration from 20 µM to 30 µM and 35 µM (Whole-population mean deformation: $D_{20\,\mu M} = 0.019 \pm 0.0001$, $D_{30\,\mu M} = 0.021 \pm 0.0001$, $D_{35\,\mu M} = 0.022 \pm 0.0001$). *p-values: 0.039, 0.032. Likewise the Young's modulus decreased significantly between the same concentrations (Whole population mean: $YM_{20\,\mu M} = 2.84\,kPa \pm 0.03\,kPa$, $YM_{30\,\mu M} = 2.36\,kPa \pm 0.02\,kPa$, $YM_{35\,\mu M} = 2.15\,kPa \pm 0.02\,kPa$). *p-values: 0.043, 0.019. Statistical significance was analyzed using a linear mixed model (R-lme4) as integrated in Shape-Out (version 2.10.0) yielding ANOVA p-values. Error values correspond to the standard error of the mean.

#### 2.16 Figure S16: Dynamic real-time deformability cytometry of 3-arm short DNA-HMPs

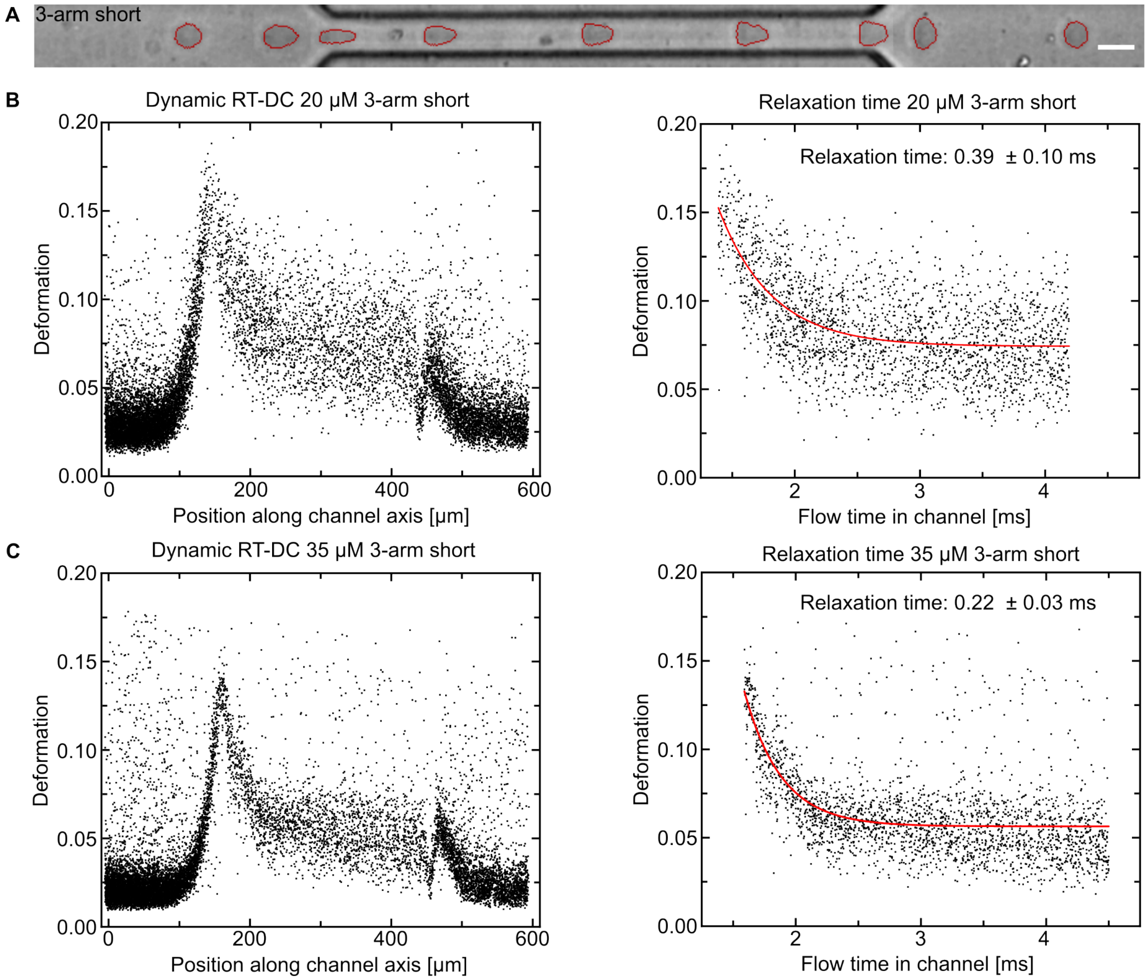

Figure S16: Dynamic real-time deformability cytometry (dRT-DC) of 3-arm short DNA-HMPs. A) Composite image of 3-arm short DNA-HMPs being deformed inside the flow channel during dRT-DC. The DNA-HMPs are spherical prior to channel-entry and initially deform strongly upon insertion into the channel. After the maximum deformation is reached, the particles relax and reach a steady-state deformation. After leaving the channel, the DNA-HMPs return to their spherical shape. Scale bar: 20 µm. B) Deformation of 20 µM DNA-HMPs plotted over the channel length during dRT-DC measurements. The DNA-HMPs initially deform strongly upon entering the channel. A steady-state of deformation is then reached within the channel following relaxation. The relaxation time of the DNA-HMPs was extracted from the decay of the exponential fit of the relaxation curve (DNA-HMP deformation over flow time). C) Deformation of 35 µM DNA-HMPs plotted over the channel length during dRT-DC measurements. The DNA-HMPs initially deform strongly upon entering the channel. A steady-state of deformation is then reached within the channel following relaxation. After the maximum deformation is reached, the particles relax and reach a steady-state of deformation. Deformation of 35 µM DNA-HMPs plotted over the flow time inside the RT-DC channel. The relaxation time of the DNA-HMPs was extracted from the decay of the exponential fit of the relaxation curve (DNA-HMP deformation over flow time). For calculation of the flow time see Experimental Section for more detail. Plotting and fitting of the data were conducted using OriginPro 2021 - Update 6 (Origin Lab Corporation).

#### 2.17 Figure S17: Dynamic real-time deformability cytometry of 6-arm flexible DNA-HMPs

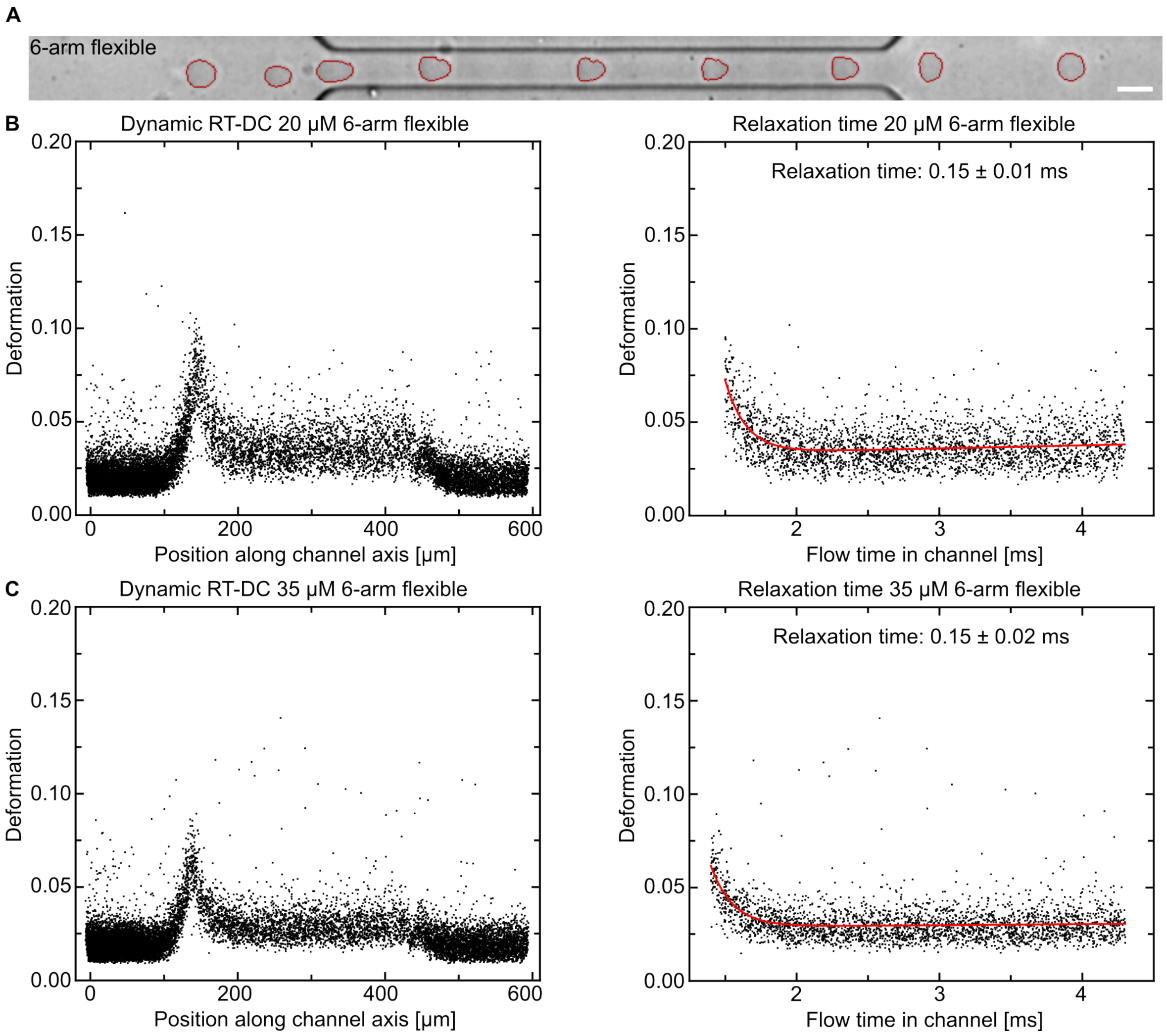

Figure S17: Analysis of 6-arm flexible DNA-HMPs using dynamic real-time deformability cytometry (dRT-DC). A) Composite image of 6-arm flexible DNA-HMPs being deformed inside the flow channel during dRT-DC. The DNA-HMPs are spherical prior to channel-entry and initially deform upon insertion into the channel, however much less than the 3-arm short DNA-HMPs. After the maximum deformation is reached, the particles relax and reach a steady-state deformation. After leaving the channel, the DNA-HMPs return to their spherical shape. Scale bar: 20 µm. B) Deformation of 6-arm flexible DNA-HMPs at 20 µM DNA concentration plotted over the channel length during dRT-DC measurements. The relaxation time of the DNA-HMPs was extracted from the decay of the exponential fit of the relaxation curve (DNA-HMP deformation over flow time). C) Deformation of 6-arm flexible DNA-HMPs at 35 µM DNA concentration plotted over the channel length during dRT-DC measurements. The relaxation time of the DNA-HMPs was extracted from the decay of the exponential fit of the relaxation curve (DNA-HMP deformation over flow time). For calculation of the flow time see Experimental Section. Plotting and fitting of the data were conducted using OriginPro 2021 - Update 6 (Origin Lab Corporation).

#### 2.18 Figure S18: Dynamic real-time deformability cytometry of 6-arm DNA-HMPs

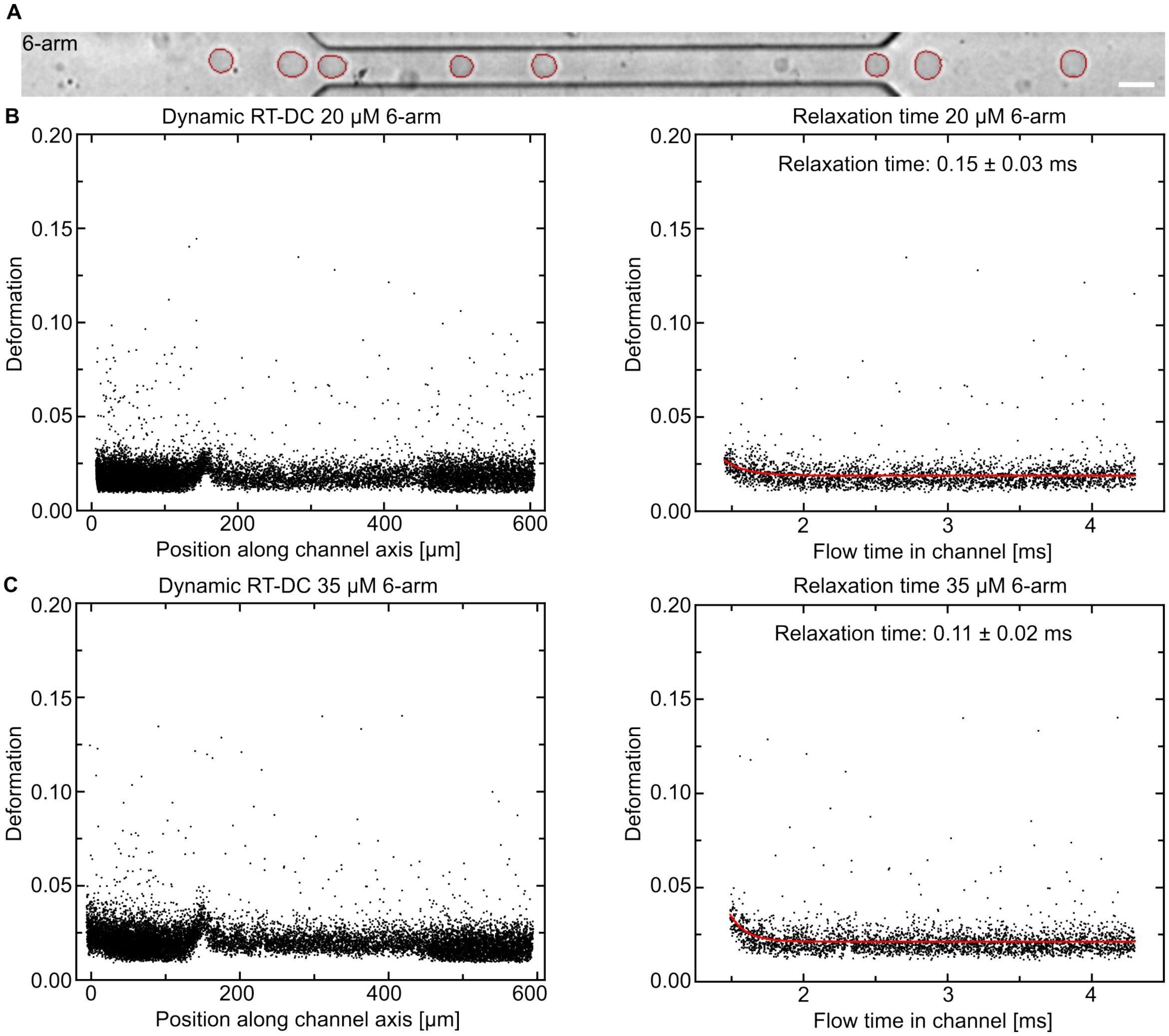

Figure S18: Analysis of 6-arm DNA-HMPs using dynamic real-time deformability cytometry (dRT-DC). A) Composite image of 6-arm DNA-HMPs being deformed inside the flow channel during dRT-DC. The DNA-HMPs are spherical prior to channel-entry and only deform slightly upon insertion into the channel; much less than any other DNA-HMPs showing a maximum deformation of around 0.03. After the maximum deformation is reached, the particles relax and reach an almost spherical steady-state deformation. After leaving the channel, the DNA-HMPs stay in their spherical shape. Scale bar: 20 μm. B) Deformation of 6-arm DNA-HMPs at 20 μM DNA concentration plotted over the channel length during dRT-DC measurements. The relaxation time of the DNA-HMPs was extracted from the decay of the exponential fit of the relaxation curve (DNA-HMP deformation over flow time). C) Deformation of 6-arm DNA-HMPs at 35 μM DNA concentration plotted over the channel length during dRT-DC measurements. The relaxation time of the DNA-HMPs was extracted from the decay of the exponential fit of the relaxation curve (DNA-HMP deformation over flow time). For calculation of the flow time see Experimental Section. Plotting and fitting of the data were conducted using OriginPro 2021 - Update 6 (Origin Lab Corporation).

#### 2.19 Figure S19: Polyacrylamide gel electrophoresis of modified and unmodified elongated linker, 6-arm linker and flexible 6-arm linker

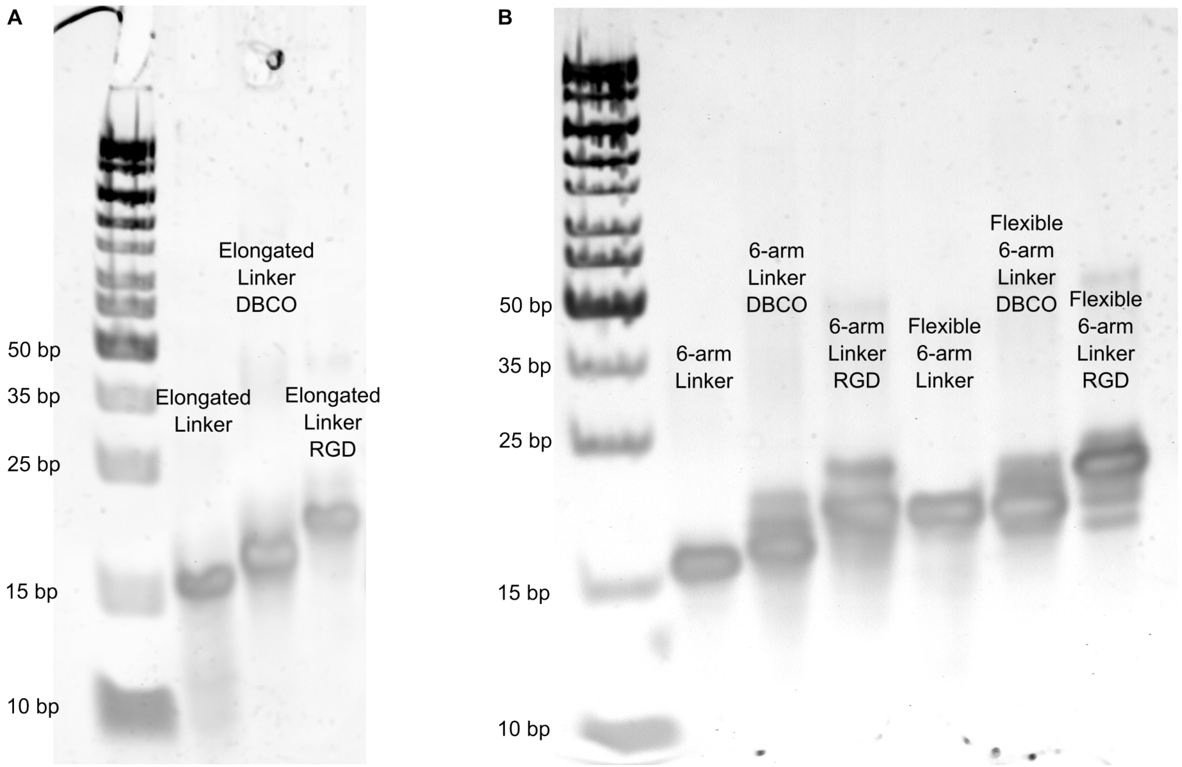

Figure S19: Polyacrylamide gel electrophoresis (PAGE) of the elongated linker, 6-arm-linker and flexible 6-arm-linker with and without modifications. A) PAGE showing the elongated linker DNA construct without any modifications, as well as the DBCO- and RGD-modified elongated linker. B) PAGE showing the 6-arm-linker and flexible 6-arm linker DNA constructs without any modifications, as well as the DBCO- and RGD-modified linkers. The marked size increase of the DBCO-modified linkers and the RGD-modified linkers above the non-modified DNA indicates the coupling of the respective modifications to the intact DNA linkers.

#### 2.20 Figure S20: Extraction of drag force amplitude via force time series

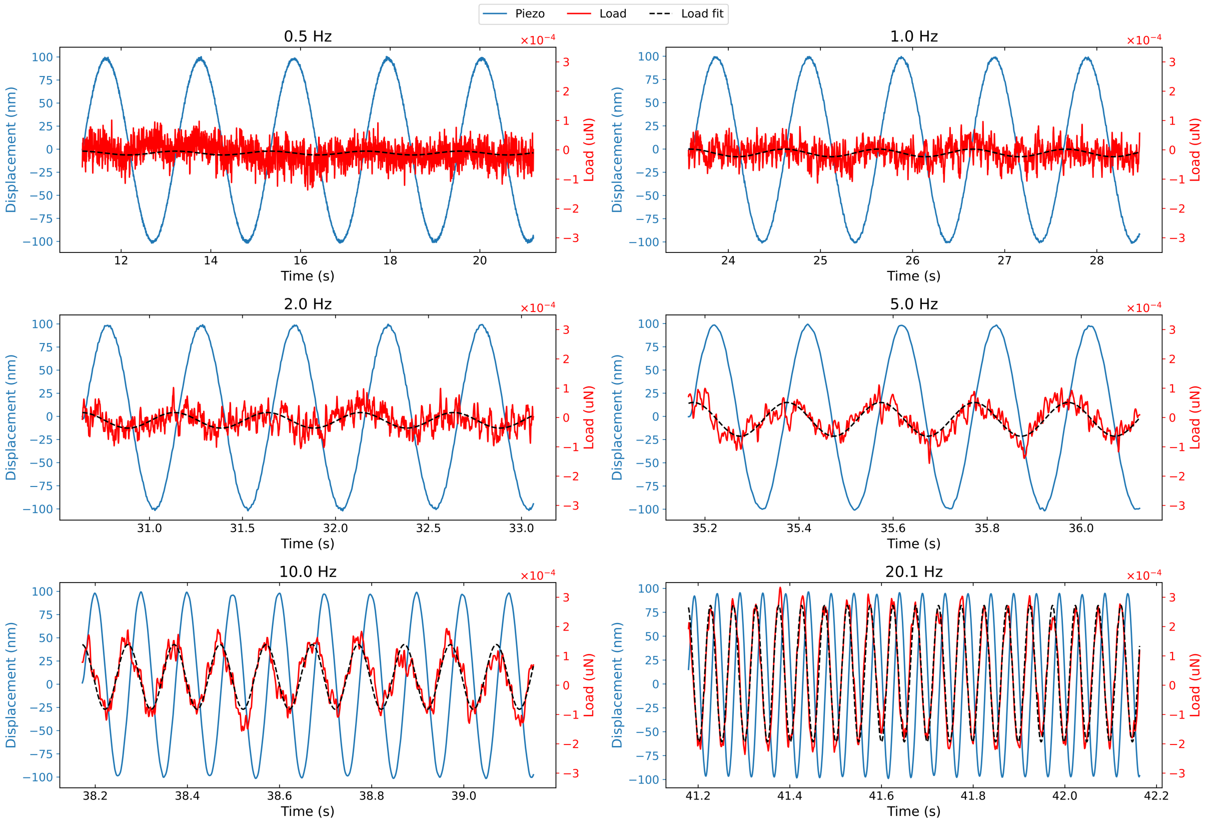

Figure S20: Drag force extraction from force time series. Force time series (red) measured by the cantilever, which is displaced in liquid by a piezoelectric actuator operating at varying frequencies. The displacement signal (blue) represents the actuator's measured position. Each subplot displays the dynamic response at different driving frequencies: 0.5 Hz, 1.0 Hz, 2.0 Hz, 5.0 Hz, 10.0 Hz, and 20.1 Hz. The black dashed line shows the sine-fitted load curve for each frequency, highlighting the increase in amplitude with frequency and the expected 90-degree phase delay, characteristic of a drag force.

#### 2.21 Figure S21: Correction of the loss modulus by the viscous drag contribution

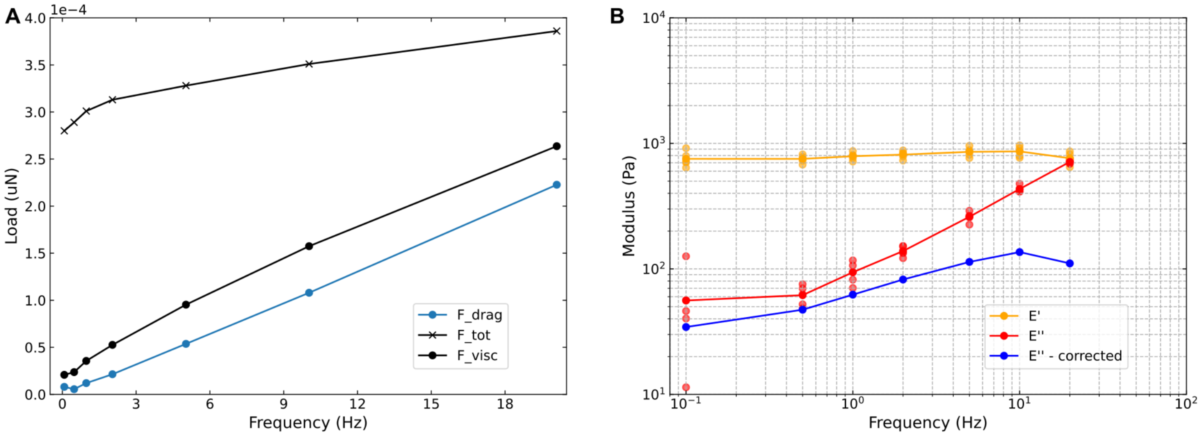

Figure S21: Correction of loss modulus E" by viscous drag. A) The load on the cantilever as a function of frequency is shown for three forces: The drag force, $F_{\mathrm{drag}}$ (blue dots), extracted as shown in Fig. S20; the total force, $F_{\mathrm{tot}}$ (crosses), provided by the manufacturer's analysis software; and the viscous force, $F_{\mathrm{visc}}$, calculated using $F_{\mathrm{tot}}$ and the loss tangent, $tan(\delta)$. The data shown represents a typical measurement for 3-arm short DNA-HMPs, where $F_{\mathrm{drag}}$ accounts for the majority of $F_{\mathrm{visc}}$. B) The storage modulus, E', and loss modulus E" are plotted against frequency for n = 5 measurements of 3-arm short DNA-HMPs. After accounting for the contribution of $F_{\mathrm{drag}}$ to $F_{\mathrm{visc}}$, a correction is made to E" (see Supplementary Note 1).

### 3 Supporting Videos

#### 3.1 Video S1: Droplet-templated formation of 3-arm short DNA-HMP over time

Overlay of confocal microscopy ($\lambda_{ex}$ = 561 nm, Cy3-labeled DNA, yellow) and brightfield time-lapse of a 3-arm short DNA-HMP forming inside a water-in-oil droplet over the course of two hours. The DNA condenses across the whole water-in-oil droplet to form a single DNA-HMP. Scale bar: 10 µm.

#### 3.2 Video S2: Fluorescence recovery after photobleaching of released DNA-HMP

Confocal microscopy ($\lambda_{ex}$ = 561 nm, Cy3-labeled DNA, yellow) time-lapse of a released 3-arm short DNA-HMP during a fluorescence recovery after photobleaching (FRAP) experiment. The bleached area does not recover after bleaching, confirming the formation of a stable gel-phased structure. Scale bar: 10 µm.

#### 3.3 Video S3: Integration of 3-arm DNA-HMPs into 3D fibroblast spheroids

Overlay of a confocal microscopy ($\lambda_{ex}$ = 561 nm, td-tomato-labeled mouse liver fibroblasts, yellow, $\lambda_{ex}$ = 405 nm, ATTO-390-labeled DNA, cyan) z-stack of 3-arm DNA-HMPs embedded into a fibroblast spheroid after 48 h of hanging drop co-culture. Scale bar: 100 µm.

#### 3.4 Video S4: Integration of 4-arm DNA-HMPs into 3D fibroblast spheroids

Overlay of a confocal microscopy ($\lambda_{ex}$ = 561 nm, td-tomato-labeled mouse liver fibroblasts, yellow, $\lambda_{ex}$ = 488 nm, ATTO-488-labeled DNA, green) z-stack of 4-arm DNA-HMPs embedded into a fibroblast spheroid after 48 h of hanging drop co-culture. Scale bar: 100 µm.

#### 3.5 Video S5: Integration of 6-arm flexible DNA-HMPs into 3D fibroblast spheroids

Overlay of a confocal microscopy ($\lambda_{ex}$ = 561 nm, td-tomato-labeled mouse liver fibroblasts, yellow, $\lambda_{ex}$ = 640 nm, ATTO-647-labeled DNA, red) z-stack of 6-arm flexible DNA-HMPs embedded into a fibroblast spheroid after 48 h of hanging drop co-culture. In some cases the fibroblasts invaded into 6-arm flexible DNA-HMPs, effectively breaking them open. We attribute this to the combination of high stiffness and more porous nature of this design. Scale bar: 100 µm.

#### 3.6 Video S6: Integration of 6-arm DNA-HMPs into 3D fibroblast spheroids

Overlay of a confocal microscopy ($\lambda_{ex}$ = 561 nm, td-tomato-labeled mouse liver fibroblasts, yellow, $\lambda_{ex}$ = 640 nm, ATTO-647-labeled DNA, red) z-stack of 6-arm DNA-HMPs embedded into a fibroblast spheroid after 48 h of hanging drop co-culture. Scale bar: 100 µm.

#### 3.7 Video S7: Deformation of a 3-arm DNA-HMP in a 3D fibroblast spheroid over time

Overlay of a confocal microscopy ($\lambda_{ex}$ = 561 nm, td-tomato-labeled mouse liver fibroblasts, yellow, $\lambda_{ex}$ = 405 nm, ATTO-390-labeled DNA, cyan) time-lapse of a 3-arm DNA-HMP being deformed in a fibroblast spheroid by the surrounding cells over the course of 13.5 h. Scale bar: 20 µm.

##### 3.8 Video S8: Deformation of a 4-arm DNA-HMP in a 3D fibroblast spheroid over time

Overlay of a confocal microscopy ($\lambda_{ex} = 561$ nm, td-tomato-labeled mouse liver fibroblasts, yellow, $\lambda_{ex} = 488$ nm, ATTO-488-labeled DNA, green) time-lapse of a 4-arm DNA-HMP being deformed in a fibroblast spheroid by the surrounding cells over the course of 14 h. Scale bar: 20 µm.

##### 3.9 Video S9: Deformation of a 6-arm flexible DNA-HMP in a 3D fibroblast spheroid over time

Overlay of confocal microscopy ($\lambda_{ex} = 561$ nm, td-tomato-labeled mouse liver fibroblasts, yellow, $\lambda_{ex} = 640$ nm, ATTO-647-labeled DNA, red) time-lapse of a 6-arm flexible DNA-HMP being deformed in a fibroblast spheroid by the surrounding cells over the course of 13.5 h. Scale bar: 20 µm.

##### 3.10 Video S10: Deformation of a 6-arm DNA-HMP in a 3D fibroblast spheroid over time

Overlay of a confocal microscopy ($\lambda_{ex} = 561$ nm, td-tomato-labeled mouse liver fibroblasts, yellow, $\lambda_{ex} = 640$ nm, ATTO-647-labeled DNA, red) time-lapse of a 6-arm DNA-HMP being deformed in a fibroblast spheroid by the surrounding cells over the course of 13.5 h. Scale bar: 20 µm.

#### 4 Supplementary Note 1: Correction for drag force

When a cantilever oscillates in a viscous medium, it experiences a force even in the absence of physical contact with a surface. This force is due to hydrodynamic drag, which depends on the viscosity and density of the medium, the size of the cantilever, and the frequency of oscillation. Thus, it needs to be determined for any given set of experimental parameters. If not accounted for, this viscous drag can be mistakenly attributed to the properties of the material being tested during dynamic mechanical measurements. To account for the viscous drag in our measurements, we applied the following steps:

**Measurement of the drag force:** We measured the viscous drag force during DMA as described in the experimental section *Measurement of hydrodynamic drag forces during microindentation*. The force amplitude at each frequency of the freely oscillating cantilever was then determined by fitting a sine wave to the data (Figure S20). Based on five individual measurements we calculated an average drag force $F_{\text{drag}}$.

**Calculating the viscous force:** Using the analysis data extracted from the original DMA measurements from the analysis software DataViewer (V2.5.0, Optics11Life), the loss tangent $tan(\delta)$ is calculated as $E''/E'$. Together with the detected load amplitude $A$, this is taken to calculate $F_{\text{visc}} = A * sin(\delta)$ (Figure S21A).

**Calculating geometrical conversion factor:** The analysis software applies a geometrical conversion factor, to convert the force to a modulus, which we back-calculate as: $C = E''/F_{\text{visc}}$.

**Correcting the loss modulus:** Finally, we correct the loss modulus $E''$ by subtracting the drag force contribution as measured earlier via $E''_{\text{corrected}} = E'' - C * F_{\text{drag}}$, which can also be written as $E''_{\text{corrected}} = E''(1 - F_{\text{drag}}/F_{\text{visc}})$.
